# Monitoring systems for resistance to plant protection products across the world: Between redundancy and complementarity

**DOI:** 10.1101/2020.07.30.228239

**Authors:** The Reflection and Research Ring on Pesticide Resistance (R4P) is constituted of, Benoit Barrès, Marie-France Corio-Costet, Danièle Debieu, Christophe Délye, Sabine Fillinger, Bertrand Gauffre, Jacques Grosman, Mourad Hannachi, Pauline de Jerphanion, Gaëlle Le Goff, Christophe Plantamp, Myriam Siegwart, Anne-Sophie Walker, Lise Nistrup-Jørgensen

**Author notes:** Invited guest of R4P to contribute to this paper. **Corresponding author:** Anne-Sophie Walker, Université Paris-Saclay, INRAE, AgroParisTech, UMR BIOGER, 78850 Thiverval-Grignon, France.

## Abstract

**BACKGROUND:** Monitoring resistance to Plant Protection Products (PPPs) is crucial for understanding the evolution of resistances in bioagressors, thereby allowing scientists to design sound bioagressor management strategies. Globally, resistance monitoring is implemented by a wide range of actors that fall into three distinct categories: academic, governmental, and private. The purpose of this study was to investigate worldwide diversity in PPP resistance monitoring systems, and to shed light on their different facets.

**RESULTS:** A large survey involving 162 experts from 48 countries made it possible to identify and analyze 250 resistance monitoring systems. Through an in-depth analysis, the features of the different monitoring systems were identified. The main factor differentiating monitoring systems was essentially the capabilities (funding, manpower, technology, etc.) of the actors involved in each system. In most countries, and especially in those with a high Human Development Index, academic, governmental, and private monitoring systems coexist. Overall, systems focus far more on monitoring established resistances than on the detection of emerging resistances. Governmental and private resistance monitoring systems generally have considerable capacities to generate data, whereas academic resistance monitoring systems are more specialized. Governmental actors federate and enroll a wider variety of stakeholders.

**CONCLUSION:** The results show functional complementarities between the coexisting actors in countries where they coexist. We suggest PPP resistance monitoring might be enhanced if the different actors focus more on detecting emerging resistances (and associated benefits) and increase collaborative and collective efforts and transparency.

## 1. Introduction & background

Plant Protection Products (PPPs) include active ingredients or organisms used to kill, alter the development of, or mitigating the deleterious effects of plant bioagressors such as animals, pathogens, and weeds. In agriculture, PPPs can select resistant bioagressor genotypes that have the inheritable ability to survive PPP concentrations that kill or inhibit the development of sensitive genotypes of the same species. Resistance is generally selected from existing bioagressor genetic variation or from *de novo* mutations (Hawkins et al., 2019; R4P, 2016). When the proportion of resistant genotypes in one bioagressor population is high enough, resistance leads to a visible decrease in the efficacy of PPP applications in the field. This is “resistance in practice”, which is considered a side effect of PPP use. It may lead farmers to carry out additional applications of higher doses of the PPP concerned, or use other PPPs with possibly less favorable ecotoxicological profiles. Bioagressor resistance has considerable economic consequences worldwide in terms of additional PPP use, compensatory agronomic practices, food production losses, and environmental pollution (Palumbi, 2001). A recent study conducted in the United Kingdom estimated that herbicide resistance can double the economic costs of weed management (Hicks et al., 2018). Other studies concluded that weed management costs could increase by between USD 85 and USD 138.ha−1 when herbicide resistance is present (Lambert et al., 2017; Zhou et al., 2015). Therefore, managing the evolution of resistance is a major challenge in crop protection, in line with the increasing social pressure for less dependence on chemical inputs in agriculture (Jørgensen et al., 2020). This concern has been translated into law in many countries and implied the development of specific regulations (Box 1) and of dedicated PPP risk indicators (Box 2).

Efficient resistance management relies: (i) on the use of bioagressor control practices that do not resort to PPP (e.g. resistant cultivars, prophylaxis, etc.), and (ii) if needed, the judicious application of PPPs following anti-resistance strategies, namely alternating PPPs, using mixtures, applying spatial mosaic practices, or dose modulation, so as to maximize the heterogeneity of selection applied on bioagressor genotypes (Délye et al., 2013; Jørgensen et al., 2017; REX_Consortium, 2013; van den Bosch et al., 2014). While the relative efficacies of the different strategies in impeding resistance development are still debated, probably because they closely depend on bioagressor biology and resistance genetics, there is a consensus that the management of resistance at the earliest stages of its development is a prerequisite for the sustainability of PPP efficacy. Designing efficient strategies for PPP resistance management requires knowledge of the resistance status and resistance mechanisms of a given bioagressor to one or several PPPs, and therefore depends on information obtained from PPP resistance monitoring and academic research (see an example in Box 3).

PPP resistance monitoring involves regular collection of data on the occurrence, frequency, and/or location of bioagressor genotypes with phenotypic and genetic characteristics making them resistant to PPPs. Resistance monitoring is used across the world to assess and curb the incidence of resistant human pathogens (Broekmans et al., 2002; Mölstad et al., 2017; Zignol et al., 2018) or animal pathogens (Werner et al., 2018), or for food safety (Mc Nulty et al., 2016; Pruden et al., 2018). In many countries, resistance monitoring is a prerequisite for PPP authorization and a key component of the post-authorization phase, and therefore falls within the jurisdiction of governmental and regulatory authorities. It also enables forecasting of in-field PPP efficacy and makes it possible to put forward preventive measures against the development of resistance. In addition, resistance monitoring is instrumental in further adapting stakeholder strategies and in preventing crop protection impasses. Resistance monitoring helps to optimize the efficient use of PPPs, maintain their efficacy and guides commercial and R&D strategies in the plant protection product industry. Consequently, there is a market for PPP resistance monitoring data, which attracts private organizations. On a different level, preventing the use of inefficient PPPs by adequate resistance monitoring contributes to the public interest objective of limiting the overall amount of PPPs used in a country and is therefore the purview of public institutions. Moreover, PPP resistance monitoring is also of interest for academic scientists studying the evolution of bioagressors. Clearly, PPP resistance constitutes a textbook example of an evolutionary process that can be observed at contemporary scales (e.g. (Blake et al., 2018; Garnault et al., 2019; Hicks et al., 2018; Labbé et al., 2007). PPP resistance monitoring is therefore implemented by a variety of actors who fall into three main categories: academic, governmental, and private. Despite the importance and the diversity of the issues associated with PPP resistance monitoring, no quantitative or qualitative information had previously been published at a worldwide level on the organization of PPP resistance monitoring. Furthermore, the handful of studies that have investigated the organization of PPP resistance monitoring all focused on a specific topic and/or considered a limited number of countries. Examples include a public-private partnership for insecticide resistance management of the diamondback moth in Hawaii (Krell et al., 2016), a public-private consortium to study fungicide resistance in Septoria leaf blotch on wheat in Europe (Jørgensen et al., 2018), institutional organization of monitoring approaches for resistance to transgenic *Bacillus thuringiensis* (Bt)-crops in four countries (Carriere et al., 2019), and an overview on herbicide resistance management across the world with few details on monitoring (Peterson et al., 2018).

This study aimed to describe the diversity of actors and features of PPP resistance monitoring within and among countries, to explore the factors underlying this diversity, and to identify possible synergies. Our general hypothesis was that functional complementarities between the coexisting actors in a given country may explain the diversity of monitoring systems observed around the world. To test this assumption, we used a multilevel case study questionnaire methodology (Eisenhardt, 1989; Yin, 2003), and more precisely, an embedded, multiple-case research design (Yin, 2003). The most important level we considered was the Resistance Monitoring System (RMS), which is an organization with its own objectives, resources, and rationales that produces and organizes PPP resistance monitoring (**Figure 1**). An RMS produces Resistance Monitoring Information Products (RMIPs), deliverables that enable actors to visualize and track the development of resistance, whether these products are publicly shared or not. An RMS may produce several different information products. At a higher level, in a given country, the coexisting RMSs shape a National Resistance Monitoring Landscape (NRML). Based on this multilevel design, we: (i) described the diversity of RMSs around the world, and tested whether the NRML could be related to country’s Human Development Index (HDI), (ii) described similarities and differences regarding the various RMSs, and (iii) characterized the interplay between actors that implement different RMSs.

**Figure 1:**
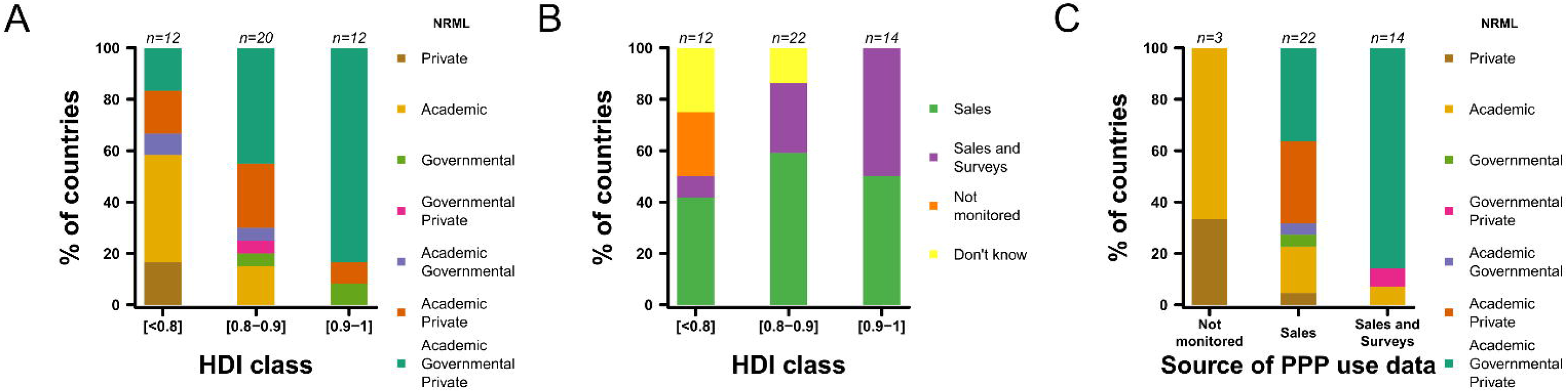
The three levels of investigation: the National Resistance Monitoring Landscape (NRML), the Resistance Monitoring System (RMS), and the Resistance Monitoring Information Product (RMIP).

## 2. Methods

### 2.1. Questionnaire design

Based on a preliminary investigation, we defined three categories of RMS, based on the type and funding of the organization in charge: Academic, Governmental, and Private. An Academic RMS is a monitoring initiative carried out by academic actors, where the main aim is the scientific study of resistance evolution. A Governmental RMS is an initiative aimed at producing publicly available information for agricultural extension services and state services for PPP evaluation and authorization, and to address PPP regulation questions (e.g. non-intentional effects of PPPs such as the selection for resistances). A Private RMS is a resistance monitoring initiative carried out by private actors, where the main aim is to detect and track the spread of resistance, with the aim of supporting R&D and PPP stewardship, as well as PPP evaluation by the authorities.

The questionnaire was composed of a series of 75 directive and 35 non-directive questions (Supporting Information 3). Both directive and non-directive questions explored the reasons and rationales of the RMS by addressing various features (informant details, country, etc.). The respondents were asked to list all the existing RMSs they were aware of in their country, as well as the reasons and rationales for each of them. One respondent could describe up to one RMS belonging to each category (maximum 3 RMS per respondent) to focus on the RMSs they know best and to avoid bias such as the overrepresentation of some countries and a strong heterogeneity in responses among respondents. The questionnaire also asked for details on each RMS listed by the respondent (e.g. category, objectives, spatial and temporal scales, methodology, funding, links with registration and regulation, communication of results, etc.). The last part of the questionnaire explored the collection and availability of data on PPP use throughout the respondents’ countries, (through sales data and/or survey). These data are useful for resistance monitoring in order to select PPPs of interest and to focus resources on regions where selection pressure toward resistance is high (see Box 2 for a detailed example). When the question was directive (multiple-choice), a free response insert was available to add an option or make a comment. Hence, several of these responses and options were re-encoded for the analysis to create groups, when relevant.

### 2.2. Data collection

The experts contacted in a preliminary phase were chosen for their ability to identify the PPP resistance monitoring initiatives and/or their own participation in such monitoring programs in their country. In addition, we tried to balance the number of experts in each major category of pesticides by sending the questionnaire to a similar number of experts in insecticide, fungicide and herbicide resistance. In a second phase, we used a snowball sampling technique: the initial respondents were invited to recruit new respondents meeting the eligibility criteria mentioned above. This sampling technique was used because of the difficulty in identifying resistance monitoring experts internationally (Heckathorn, 1997). The questionnaire was sent via a *Sphinx* software internet link from October 3, 2016 to November 22, 2017.

### 2.3. Data analysis

For each country, responses concerning RMSs were aggregated to generate comprehensive information on its NRML. Countries with one or few respondents were not excluded from the analysis. However, this criterion was taken into account when interpreting the data and comparing countries. For countries with several respondents, such as European Union countries, Australia and the United States, answers about the NRML were consistent among respondents. Similarly, for each RMS described, we asked which resistance topics were monitored (one topic = one triplet: crop × bioagressor × PPP active ingredient). These topics were then grouped according to bioagressor types (fungi, insects, weeds, and combinations of these three types). These answers where compiled for each country.

To analyze whether countries with a higher development level were characterized by a more diverse NRML, we used the 2016 Human Development Index (HDI (UNDP, 2016)), a composite statistical value of life expectancy, education, and per capita income indicators (Anand & Sen, 1994)). The influence of the HDI on the presence of each RMS category and the influence of the HDI on the type of collection of PPP use data in each country were tested using generalized linear regressions with a binomial distribution of errors and logit link function.

The effect of the category of RMS on the number of actor types participating in the choice of the monitored topics, or in the resistance data analysis, were tested using a non-parametric Kruskal– Wallis rank-sum test. All other tests of relationships between different qualitative variables were conducted by chi-squared (*χ*^2^) tests of independence. When the number of replicates was too small for a *χ*^2^ test, we used a *χ*^2^ test with Monte-Carlo simulations using 2,000 replicates to compute *p*-values, or opted for a qualitative comparative analysis (Greckhamer et al., 2008; Livne-Tarandach et al., 2015). All statistical analyses were carried out using the statistical software R 3.6.1 (R Core Team, 2019). The data and code are available in a Zenodo online repository (http://doi.org/10.5281/zenodo.3723898).

## 3. Results and discussion

The questionnaire was sent to 554 experts on PPP resistance monitoring from 74 countries, including 283 experts in Europe and 271 outside Europe. A total of 162 experts (29 %) from 48 countries (65 %) responded to the questionnaire. The countries with the highest number of respondents were mainly located in Europe and North America (**Figure 2A**). Europeans represented 64 % of the respondents (103 experts), despite our efforts to collect data from a wide range of countries. The main group of respondents was academic experts (66 % of the respondents). The other groups identified were executives in government organizations (20 %), experts from private companies (10 %), and extension services and agricultural consulting companies (4 %). The area of expertise of most respondents was herbicide or fungicide resistance (32 % and 30 %, respectively), followed by insecticide resistance (18 %), and generalist experts (more than one area of expertise, representing 20 %) (**Figure 2B**). With our questionnaire approach, we inventoried 250 RMSs worldwide, relatively well balanced among countries and among the categories defined for this study, i.e Academic, Governmental, and Private (**Figures 2C and 2D**).

**Figure 2:**
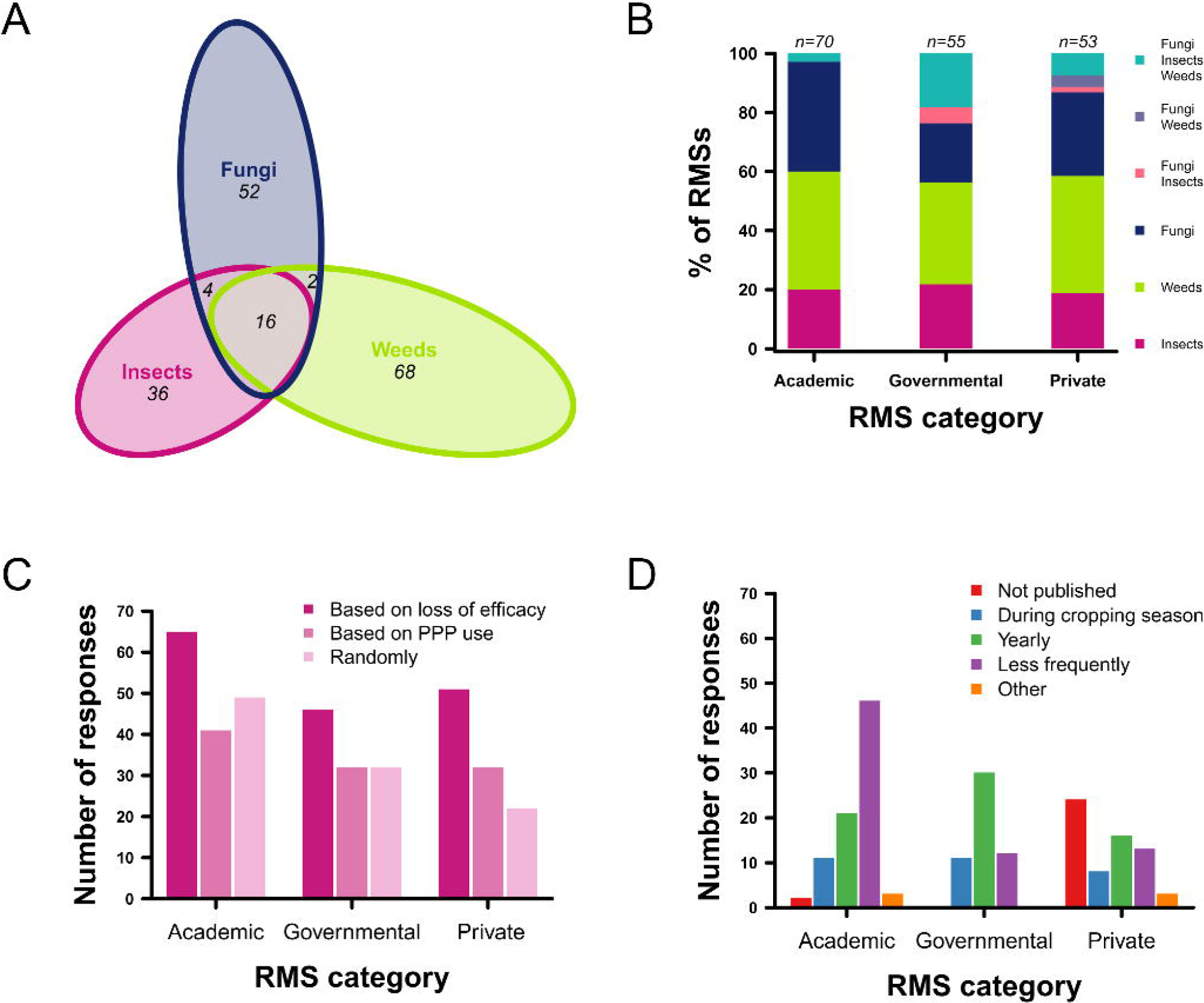
Profile of questionnaire responses: number of questionnaire respondents (A) across the world, and (B) according to their area of expertise in PPP resistance by bioagressor type. (C) Number of RMSs per RMS category described by the respondents (one respondent could describe up to one RMS belonging to each category), and (D) number of countries with at least one described RMS for each category of RMS.

### 3.1. Diversity of NRMLs around the world

#### 3.1.1. All categories of RMSs coexist in most of the respondent countries

The most frequent situation (44 % of the countries surveyed) was the coexistence of the three categories of RMS (namely Academic, Governmental, and Private). The next two most frequent situations were the existence of Academic RMSs only and the coexistence of Academic and Private RMSs (17 % of the countries surveyed each). All other situations represented less than 4 % each. The NRML was reported to be coexistence between Governmental and Private RMSs in a single country, with one respondent. Four other countries reported no RMS. However, it should be noted that for each of these five cases, the information was provided by a single respondent, thereby limiting its reliability.

#### 3.1.2. The HDI value influences NRML diversity and collection of PPP use data

The three categories of RMS were present in 83 % of the countries with an HDI > 0.9, in 45 % of the countries with an HDI between 0.8 and 0.9, and in 17 % of the countries with an HDI < 0.8 (**Figure 3A**). Importantly, countries with an HDI < 0.8 had the least diversified NRMLs, with an overrepresentation of Academic RMSs (10 out of 12 had Academic RMSs, including five with only Academic RMSs). Consistently, the presence of Governmental RMSs was significantly more likely in countries with a high HDI (*p* = 0.018, **Figure S1**.**1**, supporting information). By contrast, the occurrence of Academic and Private RMSs did not significantly correlate with HDI (*p* = 0.429 and 0.746, respectively, **Figure S1**.**1**, supporting information).

**Figure 3:**
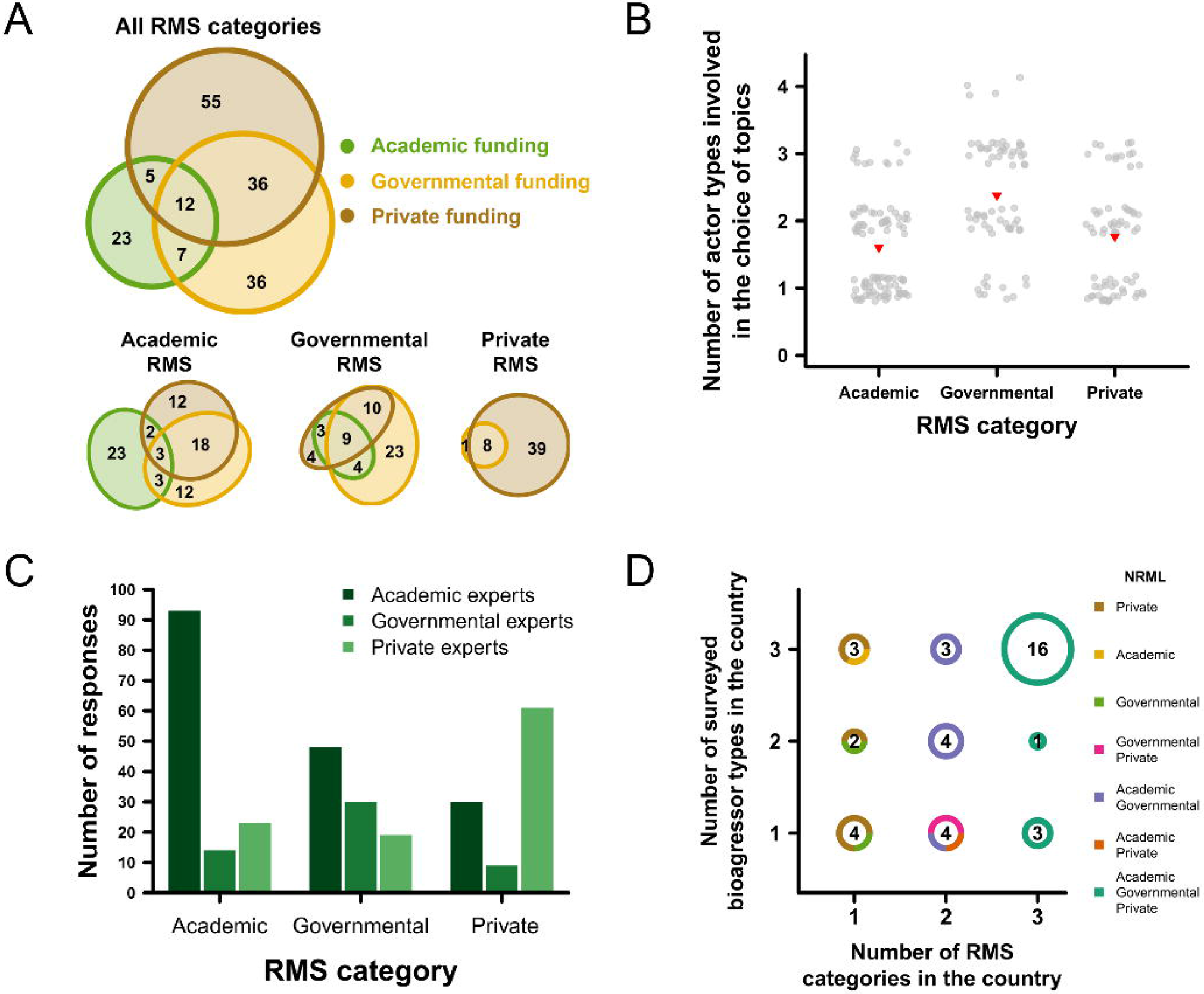
Resistance monitoring at the country level: (A) relationship between the Human Development Index (HDI) of the countries surveyed and the composition of their National Resistance Monitoring Landscape (NRML), (B) relationship between the HDI and the source of data on PPP use, (C) relationship between the sources of data on PPP use in the countries surveyed and their NRML.

PPP use was monitored through records of sales data from PPP distributors (53 %), through consumer surveys on PPP use (8 %), or through a combination of both (23 %). Data on PPP sales were collected mainly at the national and regional scales (47 % and 33 %, respectively), while surveys on PPP use were conducted at more local geographic scales (22 % at the field level, 42 % at a regional scale, and 17 % at the national scale). The procedure to collect data on PPP use differed according to the HDI: the higher the HDI, the higher the proportion of countries using both sales and surveys to collect data on PPP use (**Figure 3B**). Consistently, countries with high HDIs were more likely to use data on PPP based on ‘sales’ and ‘survey data’ to adapt their monitoring systems (*p* = 0.019 and 0.058, respectively, **Figure S1**.**2**, supporting information).

Interestingly, the more comprehensive the collection of data on PPP use, the more diverse the NRML (**Figure 3C**, *χ*^2^ = 10.6, Monte-Carlo *p* = 0.008). In the three countries where no collection of data on PPP use was reported, the NRML relied on only one category of RMS. Conversely, 87 % of the countries monitoring PPP use through both sales and surveys had the three RMS categories. This association may be found primarily in countries where governmental policies are being implemented, in particular to mitigate the environmental impact of agriculture.

### 3.2. Features of RMSs

#### 3.2.1. Governmental and Private RMSs are generalists, while Academic RMSs are more specialized

Regardless of their category, 46 % of the RMSs considered a maximum of five topics per year and 43 % considered from 5 to 20 topics per year. Only 11 % of the RMSs managed more than 20 topics per year. Private RMSs were the most involved in managing a high number of topics (47 % of the RMSs surveying more than 20 topics were Private).

Among the RMSs identified, 38 % produced RMIPs on herbicides, 29 % on fungicides, 20 % on insecticides, and 12 % on at least two bioagressor types (**Figure 4A**). Most RMSs were dedicated to a single bioagressor type, regardless of the RMS category. The proportion of RMSs investigating more than one type of bioagressor was significantly higher in Governmental RMSs (24 %), compared to Private (13 %) and Academic (3 %) ones (**Figure 4B**) (*χ*^2^ = 12.3, Monte-Carlo *p* = 0.002). Governmental RMSs specialized in fungi only were proportionally less apparent. Academic RMSs appeared to be more specialized, i.e. focused on a limited number of bioagressors type. This is most likely because Academic RMSs are usually part of broader research programs aimed at understanding the mechanisms and processes involved in the selection of resistance and at predicting the evolution of resistances or the associated risks. By contrast, governmental and private actors are mostly interested in obtaining an accurate picture of the resistance status for management or regulation purposes.

**Figure 4:**
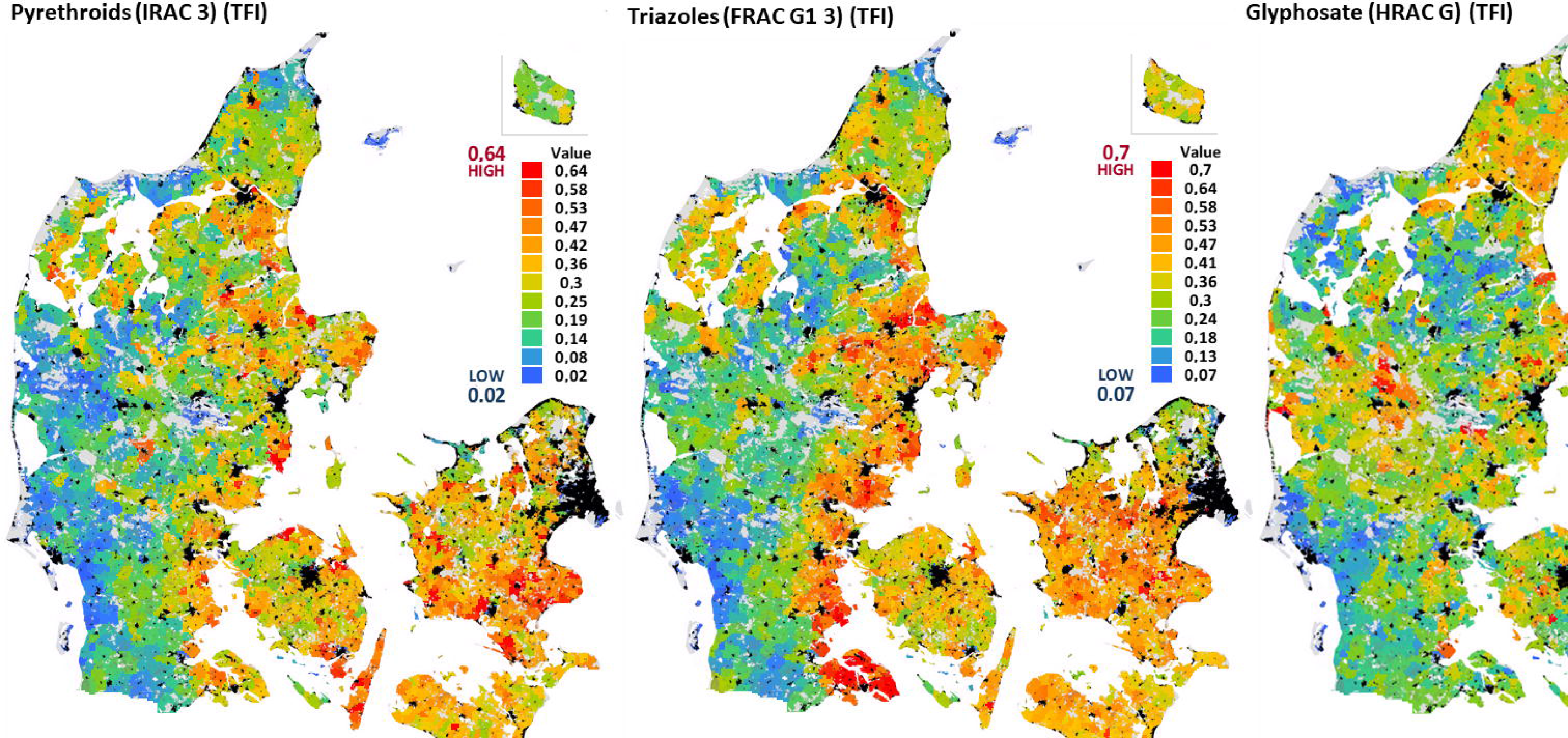
Characteristics of the different Resistance Monitoring System (RMS) categories: (A) number of RMSs and bioagressor types monitored (178 out of the 250 described RMSs included information on the type of bioagressor monitored); (B) proportions of the types of bioagressor monitored according to the RMS categories; (C) orientation of the field sampling protocol according to the RMS categories (200 out of the 250 described RMSs included information on the orientation of the sampling protocol); (D) timing and frequency of publication of Resistance Monitoring Information Products (RMIPs) according to the RMS categories (200 out of the 250 described RMSs included information on the timing of publication).

#### 3.2.2. All RMSs address emerging and established resistances, but focus mainly on loss of PPP efficacy

RMSs aim either at detecting emerging resistances or at assessing the extent of established resistances, since it can affect the type of resistance management implementation, more or less preventive. Irrespective of their category, the vast majority of RMSs had both objectives (71 %), whereas 16 % and 13 % of the RMSs focused exclusively on established resistances or emerging resistances, respectively. We did not detect a significant correlation between the number of RMS objectives and the number of RMS categories in the NRML (*χ*^2^ = 4.68, Monte-Carlo *p* = 0.319). However, for countries with a single category of RMS, the proportion of RMSs with both objectives tended to be slightly lower (60 %) than in the countries where two or three categories of RMSs coexisted (64 % or 74 %, respectively).

The sampling procedure tended to differ between RMS categories (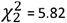, *p* = 0.054). Regardless of the RMS, the most frequent cause for field sampling was a reduction or disruption of treatment efficacy represented by 42 %, 42 %, and 49 % of responses in Academic, Governmental and Private RMSs, respectively (**Figure 4C**). Regarding the second sampling procedure, Academic and Governmental RMSs tended to use more random sampling than Private RMSs (32 % and 29 % *vs*. 21 %, re spectively), while Private RMS sampling procedures were more often based on selection pressure, i.e. on data on PPP use (31 %).

#### 3.2.3. Data collection methods and analytical techniques are similar among RMSs

Collecting relevant metadata associated with the history of the fields sampled (crop, GPS coordinates, bioagressor intensity, and history of PPP used) is an important part of resistance monitoring. These data can also be used for statistical and modelling analyses, which are necessary to assist the development of management strategies. We therefore surveyed how metadata were collected: on paper, digitally, or both. Using paper forms only was the most represented data collection methods in all RMS categories (56 %). A small trend was found for exclusively digital metadata collection that was more widespread in Academic RMSs than in the others.

Analytical techniques, including bioassays, biomolecular methods, or biochemical approaches, differ in cost, sensitivity, and specificity (R4P, 2016). There were no differences in the respective use of these different techniques for resistance detection among RMS categories (*χ*^2^ = 11.1, Monte-Carlo *p* = 0.550), despite different financial constraints between RMS categories, and despite the need for different technological expertise. Bioassay was the most frequently used technique in all RMSs (95 %), while 77 % and 34 % of the RMSs used biomolecular and biochemical tests, respectively. The use of two different analytical techniques in the same RMS was the most frequent (46 % of the RMSs), with almost exclusively a combination of bioassays and biomolecular techniques. RMSs using only one technique (24 % of the RMSs) used bioassays in the vast majority of cases (83 %). Lastly, 30 % of the RMSs, especially those investigating herbicide and insecticide resistances, combined the three techniques. Interestingly, non-target site resistance due to detoxication enzymes is more frequent in these types of bioagressors and the analysis of this resistance requires a multiscale approach from gene to phenotype (Hawkins et al., 2019).

#### 3.2.4. Diffusion and publication of the RMIPs vary according to the category of RMS

Publishing clear, up-to-date, and relevant RMIPs is important to assist growers and stakeholders in implementing efficient actions to manage resistance. RMIP publication significantly differed among RMS categories (*χ*^2^ = 50.1, Monte-Carlo *p* < 0.001, **Figure 4D**). The vast majority of the RMIPs were published (87 %). Nearly all unpublished RMIPs (92 %) were from Private RMSs. Unpublished RMIPs might be used in post-authorization reports, which are not considered formal publications and are restricted to official authorities. Alternatively, RMIPs may be considered strategic and fall under corporate proprietary information for private stakeholders and are intended for internal use only (for commercial and R&D strategies) By contrast, Governmental RMSs systematically published RMIPs, usually on an annual basis (57 %), which probably aimed at assisting resistance management and are intended to inform a broad audience. Academic RMSs published RMIPs on a less regular basis, in line with their main objective of publishing scientific articles in peer-reviewed journals, which implies combining multi-year monitoring data for population studies, resistance evolution assessments, or investigations of resistance mechanisms (e.g. Blake et al., 2018; Garnault et al., 2019; Hicks et al., 2018; Labbé et al., 2007).

### 3.3. Complementarities and interplays between RMSs and actors

#### 3.3.1. RMS funding

Private actors such as agrochemical companies or PPP retailers (grower cooperatives, vendors) were the main funders of RMSs (**Figure 5A**). They were the sole funding source for 32 % of the RMSs identified, and one of the funding sources for 30 % of the RMSs. The strong implication of private actors in RMS funding likely reflects the strategic impact of PPP resistance on their business for both regulatory obligations and their interest in PPP durability as well as to guide their strategy for the development of new active substances by identifying and understanding resistance mechanisms. All funding combinations were represented, but single-source funding dominated the picture (66 %, **Figure 5A**). However, different situations were observed according to the RMS category when considering self-funding (*χ*^2^ = 52.5, Monte-Carlo *p* < 0.001). Most Private RMSs were funded by private funds only (81 %), while private funding also supported Academic and Governmental RMSs. By contrast, Academic RMSs mixed all possible funding sources and academic funding supported almost exclusively Academic RMSs (**Figure 5A**). We suggest that the overall lower academic funding dedicated to resistance monitoring has two main causes: firstly, academic endowments are generally insufficient to support extensive or continued monitoring, and secondly, resistance monitoring is needed on a less regular basis for resistance research. Surprisingly, one out of 48 Private RMSs was entirely financed by governmental funds and 4 out of 53 Governmental RMSs were entirely financed by private funds. Overall, co-funding from two or three sources was reported for 35 % of the RMSs (28 % and 7 %, respectively). Co-funding was more frequent for Governmental RMSs (49 %) than for Academic (36 %) or Private ones (17 %).

**Figure 5:**
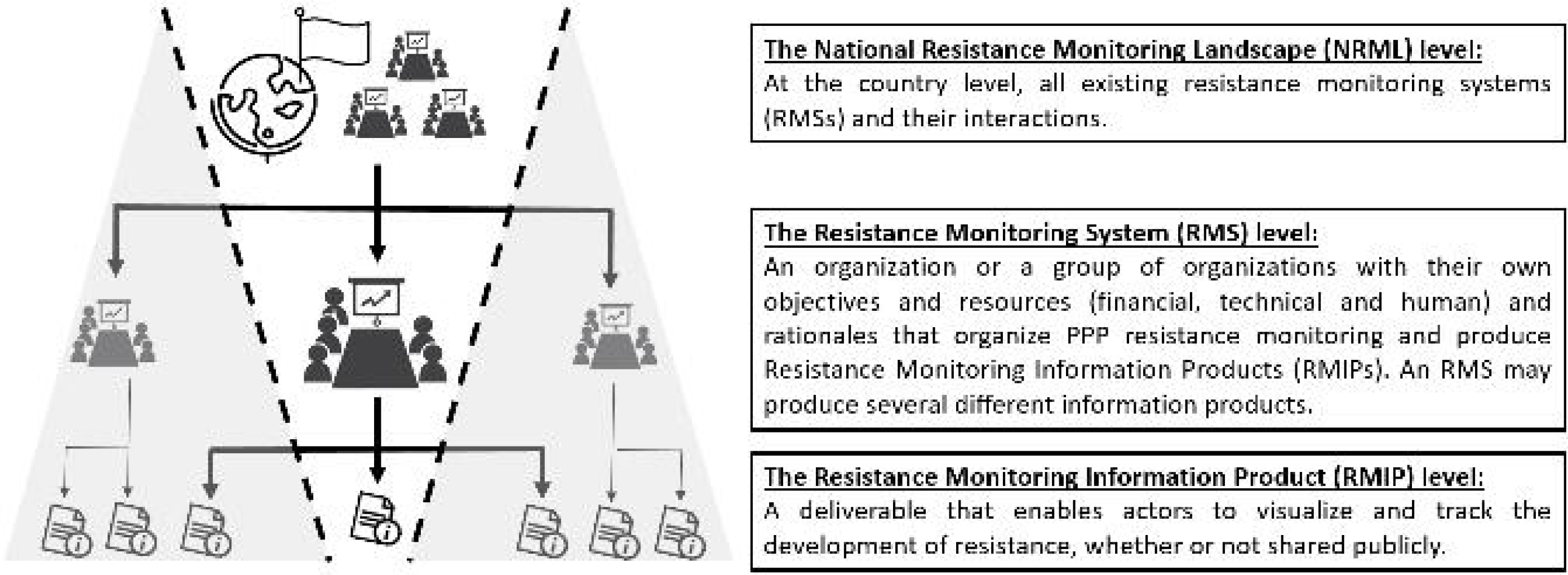
Complementarities of Resistance Monitoring Systems (RMS) and actors: (A) distribution of RMS funding sources, for all RMS categories taken together, for Academic RMSs, for Governmental RMSs, and for Private RMSs (174 out of the 250 described RMSs included information on funding); (B) number of actor types participating in the choice of topics for each RMS category (227 out of the 250 described RMSs included information on the actor types participating in the choice of topics); (C) type of experts in charge of data analysis and interpretation depending on RMS categories (219 out of the 250 described RMSs included information on the type of experts in charge of data analysis); (D) influence of the number of RMS categories in the country and the National Resistance Monitoring Landscape (NRML, in color) on the number of surveyed bioagressor types in the country. Circle size is proportionate to the number of countries (stipulated in the middle).

#### 3.3.2. Officially-standardized protocols are more conducive to public decision-making

RMIPs can be used to authorize, improve, or ban PPP use. In countries where there is official standardization of the resistance monitoring protocol (recognized as “official” by the stakeholders), changes in PPP authorization or in recommendations of PPP use is more frequent following resistance detection (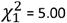, *p* = 0.025).

#### 3.3.3. Governmental actors appear to be the most able to federate and enroll a wide variety of stakeholders

The choice of the topics monitored by Governmental RMSs involved a wider range of actors (public administration, academics, companies, and others) than other RMS categories (Kruskal–Wallis 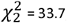, *p* < 0.001, **Figure 5B**). Twice as many partners were involved in Governmental RMSs compared to Academic and Private ones. Governmental actors lead may foster participation by others stakeholders because public involvement legitimizes the collective action.

#### 3.3.4. Academic actors are the major providers of capabilities and knowledge in the analysis and interpretation of monitoring data

Data analysis is another important aspect of PPP resistance monitoring. It requires specific knowledge adapted to the sampling design and the analysis methods used to detect and/or quantify resistance. Regardless of the RMS, the number of types of experts who analyze data is the same (ca. 1.5 persons; Kruskal–Wallis 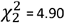, *p* = 0.080). When considering all RMSs together, academics represent more than half of the experts involved in data analysis (52 %, **Figure 5C**).

The type of experts in charge of data analysis and interpretation differed according to the RMS categories (*χ*^2^ = 50.7, Monte-Carlo *p* < 0.001). Data from Private RMSs were mostly analyzed by private actors (61 %, **Figure 5C)**, but academics were also substantially involved in the analysis of Private RMS data (30 %). The vast majority of data from Academic and Governmental RMSs was analyzed by academic actors (72 % and 50 %, respectively). Because PPP resistance data can have an impact on sales and market shares, they can be considered strategic by private stakeholders, from an economic point of view. This may explain why, despite the expertise of academics, the majority of the analyses of Private RMS data were conducted by the private stakeholders themselves. Conversely, Governmental RMSs may lack analytical skills and/or wish to avoid conflicts of interests, which may explain why they rely on academic rather than on private researchers for assistance with data analysis when external expertise is needed (DeAngelis, 2000; Robinson et al., 2013).

### 3.3.5. Coexistence of different RMSs in a country increases the diversity of the types of bioagressors surveyed

One of the greatest challenges in monitoring PPP resistances is the diversity of the topics to be considered, which implies dealing with a wide range of bioagressors, each with its own biological and genetic specificities. Eighty percent of the countries with the three categories of RMS (representing 40 % of all countries with respondents) had the three types of bioagressors surveyed (**Figure 5D**). Below three RMSs, no specific NRML composition was associated with a higher number of types of bioagressors surveyed. NRMLs were more often incomplete (not monitoring the three types of bioagressors) when only one or two categories of RMS coexisted in the country (only 33 % and 27 % monitored the three types of bioagressors, respectively). As a result, a diversified NRML seems to be correlated with more diverse and broader bioagressor monitoring.

## 4. General discussion

Monitoring PPP resistance is pivotal for implementing integrated and sustainable bioagressor management. This is the first study of this magnitude investigating the diversity of the RMSs throughout the world. Our study contains unavoidable bias, prominent among which is an over-representation of European Union and academic respondents (64 % and 66 %, respectively), while they represented 51 % and 65 % of the experts to whom the questionnaire was sent, respectively. This might be a collateral effect of our “snowball” sampling strategy, with most respondents being academics. Our work nevertheless reveals contrasted NRMLs, possibly driven by variable incentives (Box 1). We identified the (co)existence of three categories of key actors: academic, governmental, and private organizations that support three different categories of RMSs. The distinction between Academic and Governmental RMS categories could be questioned, as the two categories might overlap and usually benefit largely from public funding. However, in some countries, academic actors such as universities might be private and, beyond funding, the results of our study support our initial distinction as the three RMS categories differ in many respects, including organization, capacities, and purposes.

All three categories of RMSs generate RMIPs that differ in many ways: their degree of independence towards their funding sources, their intended use, the possibility for their public release, and their frequency of publication when published. However, the three categories of RMSs share common data collections methods and analytical techniques. They also present similarities in their objectives and sampling protocols. Interestingly, while 84 % of the RMSs claim to focus on emerging resistances, only 28 % use sampling protocols that may effectively identify emerging resistances, i.e. sampling “hot spots” based on PPP use data (Box 2). This suggests that RMSs in fact target established resistances via targeted or random samplings. This argues in favor of directing more effort toward promoting sampling protocols enabling the detection of emerging resistances, as early implementation of resistance management is most effective.

We also investigated the interplay between private, academic and governmental actors and identified a pivotal role for governmental actors. When a Governmental RMS is present in a country, especially a country with a high HDI, the diversity of types of surveyed bioagressors is higher, twice as many actor types are involved in the choice of topics, and data on PPP use are more comprehensive. This may be due to legal requirements for post-authorization procedures, a governmental prerogative. In addition, Governmental RMSs appear to be best at fostering co-funding from several actor types with half Governmental RMSs funded by at least two actors’ type. Governmental actors may be more trustworthy in the eyes of other actors. Altogether, this supports the relevance of generalized involvement of governments and/or their representatives in the monitoring systems (Fuglie & Toole, 2014; Krell et al., 2016; Wang et al., 2013). Governmental actors ability to include a broader range of partners seems to be beneficial as it combines the interests of the general public, producers, independent advisors, and markets, while promoting multi-actor synergies (Pinkse & Kolk, 2012).

Our study quantified assets that are intuitively associated with each actor, as reported in the literature (Hurley & Frisvold, 2016; Krell et al., 2016). The results suggest that private actors have a strong funding capacity and are typically active in major agricultural regions, academic actors focus on few cases and are best in data analysis capacity and scientific knowledge, and governmental actors have a major focus on disseminating information on resistance. As a result, there are obvious complementarities between the different RMS types. Resistance monitoring at a national scale could be improved if the different actors increase collaborative and collective efforts to i) enhance official standardization of the monitoring protocols, ii) focus more on emerging resistances using PPP use data to design field sampling (Box 2), iii) enhance their coordination in the choice of topics surveyed, and iv) publish transparently and regularly RMIPs. We suggest these increased collaborative efforts should eventually end up with combined academic, governmental and private capacities in joint RMSs, as put forward during the Worldwide Insecticide-resistance Network Workshop in Brazil (Corbel et al., 2017), or in a recent study on co-evolutionary governance of resistance monitoring (Jørgensen et al., 2020). Our suggestion is similar to those made recently by Carrière et al (2019) on the monitoring of insect resistance to transgenic Bt crops. The proposed type of joint RMS was not found in our survey, possibly because of the design of our questionnaire that included an *a priori* separation between Academic, Governmental and Private RMSs. However, given that the questionnaire included non-directive questions, the existence of joint RMSs would likely have emerged if it had been a frequent occurrence. Beyond potential direct synergies in joint RMSs, there are indirect synergies when Academic, Governmental and Private RMSs coexist in the same NRML (see example in Box 3). On the one hand, this cohabitation might generate redundancy in resistance topics surveyed. On the other, cohabitation increases the diversity of types of bioagressors surveyed in the country, and thereby generates mutual benefits by increasing the performance of monitoring programs. This might be due to emulation and competition among actor types, in particular for emerging resistance of significant economic interest.

Surveying changes in national RMSs and NRMLs over time should make it possible to identify additional complementarities, synergies, or tensions between RMSs.For example, Private and Governmental RMSs may clearly have antagonistic interests regarding data publication or transparency. Additional, more focused investigations could also reveal more complex NRMLs than inventoried here, in particular for regions of the world that are less represented in our survey, such as Africa or Oceania. Thanks to the present paper, we may expect more respondents from countries outside EU in future surveys. Importantly, our study investigated only RMSs and the production of RMIPs. A step forward would be to include the effective impact of the RMIPs (e.g., to adapt strategies for PPP use) on their final recipients, with each recipient category acting at their own geographical scale (e.g., growers, salespersons, technical consultants, and scientists). It would be interesting to identify the category or type of organization of RMSs that effectively prompts the final recipients to adapt or change their practices on the basis of the RMIPs. The highest utility of an RMS is no doubt reached when farmers effectively take into account the information provided by RMIPs when designing their bioagressor control tactics (Givens et al., 2011; Johnson & Gibson, 2006; Leach et al., 2019; Llewellyn et al., 2007; Ulber & Rissel, 2018). The potential impact of RMIPs depends on four characteristics: transparency, pedagogy, regular updates, and broad diffusion (via extension services, agricultural newspapers, and journals or websites). National and international groups that review and diffuse RMIPs could be key actors alongside RMSs (see examples in SI 2).

Our survey described a large number of features, but did not evaluate RMS quality. RMS quality may vary depending on financial constraints, on the expertise of the actors involved, and on the operational constraints related to the bioagressors monitored (Ambec & Desquilbet, 2012; Guedes, 2017). Based on our results, approaches could be implemented to evaluate RMS quality with regard to their objectives, and assess the quality of their output RMIPs. One such approach is the OASIS method (Hendrikx et al., 2011), which enables an in-depth analysis of the implementation and the quality of an epidemiological surveillance system, and the identification of recommendations for improvement.

## 5. Conclusion

Studying the worldwide diversity of organizations monitoring PPP resistance was a methodological challenge, and our study, although explorative and not without some unavoidable bias, is therefore an innovative and seminal research on an unexplored aspect of this crucial global issue. Charles Darwin wrote “In the long history of humankind (and animal kind, too), those who learned to collaborate and improvise most effectively have prevailed” (Darwin, 1859). Our survey revealed a diversity of RMSs with, in most countries, different categories of RMSs coexisting. Why and how could this “functional redundancy” be maintained? And why did the most effective RMS not prevail and lead the others to disappear? An analysis of the distinctive resources and the related outcome advantages of each RMS category suggests that better than mere coexistence, there is complementarity among them. The overall efficiency of an NRML could be improved if the different RMSs in different categories were merged, or better, moved towards open and transparent collaboration, and the different actors’ capabilities were pooled. This is where our study joins Darwin’s quotation. However, the benefits of RMS collaboration across categories have only been broached by the results of our explorative study, and need to be tested and studied more closely.

The antagonisms and synergisms between the different types of monitoring need to be investigated more in depth, as well as the expected benefits of collaboration. As a result, templates for multi-actor structures may be proposed to build efficient and comprehensive RMSs generating quality data and having a tangible impact on agricultural practices.

## 6. Acknowledgments

We would like to thank all the respondents who kindly participated in this survey. The original idea for this study came from a meeting of the EPPO ‘resistance panel’. This project was partly funded and supported by the French Agency for Food, Environmental and Occupational Health & Safety (ANSES) via the tax on sales of plant protection products. The proceeds of this tax are assigned to ANSES to finance the establishment of the system for monitoring the adverse effects of plant protection products, called ‘phytopharmacovigilance’ (PPV), established by the French Act on the future of agriculture dated October 13, 2014. The R4P network is supported by the Plant Health division of INRAE.

## 7. Glossary

**Bioagressor:** a living organism detrimental to crop production that can be an animal (arthropod, rodent, etc.), a plant (weed), or a phytopathogenic microorganism (bacterium, fungus, etc.).

**Emerging resistance:** early phase of resistance evolution, when resistant bioagressor genotypes are present at very low frequencies in one bioagressor population.

**Established resistance:** later phase of resistance evolution, when resistance disrupts bioagressor control in the field because the frequency of resistant bioagressor genotypes in the bioagressor population is sufficiently high.

**HDI**: Human Development Index, a composite statistical value of life expectancy, education, and per capita income indicators.

**Hot spot**: any area or place of known high levels of diversity or quantity of PPP use.

**NRML:** National Resistance Monitoring Landscape. At the country level, all existing resistance monitoring systems (RMSs) and their interactions.

**PPP:** Plant Protection Product. It includes active ingredients or organisms used to kill, alter development of, or mitigate the deleterious effects of plant bioagressors (animals, pathogens, weeds).

**PPP Resistance:** (i) Natural, inheritable ability of a bioagressor genotype (mutant, resistant genotype) to survive PPP concentrations that kill or inhibit the development of wild-type genotypes of the same species (sensitive genotypes); (ii) outcome of the adaptive evolution of bioagressors as a result of the selection pressure for less-PPP-sensitive genotypes exerted by PPPs.

**Resistant genotype:** bioagressor genotype having PPP resistance.

**Resistance monitoring:** regular collection of data on the occurrence, frequency, and/or location of resistant bioagressor genotypes.

**Resistance topic:** Crop × Bioagressor x PPP.

**RMIP**: Resistance Monitoring Information Product. A deliverable that enables actors to visualize and track the development of resistance, whether or not shared publicly.

**RMS:** Resistance Monitoring System. An organization or a group of organizations with their own objectives and resources (financial, technical and human) and rationales that organize PPP resistance monitoring and produce Resistance Monitoring Information Products (RMIPs). An RMS may produce several different information products. In this study, we defined three categories of RMS, based on the type of organization and their funding, namely

**--Academic RMS:** a resistance monitoring initiative carried out by academic actors, where the main aim is the scientific study of evolution of PPP resistance of bioagressors.

**--Governmental RMS:** a resistance monitoring initiative carried out by governmental actors, aimed at producing publicly available information for agricultural extension services and state services for PPP evaluation and authorization, and to address PPP regulation questions (e.g. non-intentional effects of PPPs such as the selection for resistances).

**--Joint RMS:** an ideal RMS that combines several types of actors (academic, governmental or/and private) managing the objectives and resources of resistance monitoring together.

**--Private RMS:** a resistance monitoring initiative carried out by private actors, where the main aim is to detect and track the spread of resistance, with the aim of supporting R&D and PPP stewardship, as well as PPP evaluation by the authorities.

## 8. Boxes

### Box 1

**Rationales for resistance monitoring across the world**

NRMLs with the three RMS categories were more frequent in countries in the highest HDI classes (see Figure 3A). Beyond important differences in national wealth, one reason for this unequal pattern of resistance monitoring organization is that countries may have different incentives when authorizing PPPs. Safety for users and the environment is a common basic incentive for all countries. Efficacy is another crucial incentive when authorizing PPPs. Some countries consider that resistance-concerned PPPs will mechanically disappear from the market because of decreasing efficacy (e.g. resistance risk assessment is not a requirement in the US Pesticide Registration Improvement Extension Act PRIA 4, nor in New Zealand or Japan (resources in SI2)). In this way, it is considered a company responsibility to anticipate the resistance risk, to market sustainable products, and to recommend good practices. Other countries are more concerned about the selection of resistance and its impact on PPP efficacy. In fact, the rapid build-up of resistance that led to historical dramatic loss of efficacy in some situations (e.g. resistance to benzimidazoles or QoIs in many pathogens, resistance to diamide insecticides in Lepidopteran pests, resistance to ACCase inhibitors in grass weeds) certainly helped convince authorities of the importance of resistance prevention and management strategies to support farmers with appropriate information on resistance. The social pressure towards an agricultural model that is less dependent on chemical inputs may also have strengthened this concern, as resistance management implies substituting PPPs by alternative control methods and prevents inefficient spraying. This has been translated into law, especially in the European Union (e.g. Directive 2009/128/EC, (Parliament, 2009)), with the development of specific regulations (e.g. the “plan Écophyto” in France, which promotes integrated bioagressor management and non-chemical bioagressor control to reduce pesticide consumption). Resistance risk assessment is also a recommendation of the Food and Agriculture Organization of the United Nations (FAO) in its guidelines for the authorization of PPPs (resources in SI2). The requirement for resistance management is included in authorization dossiers in Australia (resources in SI2) and in EU countries. In fact, the systematic evaluation of resistance in marketing authorization dossiers for PPPs has been carried out in some EU countries for years, before EU regulations came into force. Then, resistance risk assessment was generalized at the European level in 1993 via Directive 91/414/EEC, then in 2011 via Regulation (EC) No. 1107/2009, which is applied by all EU members states as a whole. The pre-marketing assessment makes it possible to take into account resistance phenomena and to anticipate problems, in particular concerning certain active ingredients that are subject to a high risk of resistance development. Some limitations on PPP use can be issued after evaluation of the marketing authorization application. Dossier requirements include sensitivity baselines, potential risk of target resistance development, a cross-resistance analysis, population status for resistance and efficacy, and recommendations for resistance management. Similarly, China recently levelled up its requirements on resistance risk assessment (Regulation on the Administration of Pesticides – Decree 677, resources in SI2), which in time may change the overall picture of resistance monitoring internationally.

### Box 2

**Spray intensity maps provide potential information on hot spots for resistance development**

It is generally accepted that the higher the spray intensity with PPPs with the same mode of action, the higher the risk of resistance selection. Based on this, resistance monitoring would be focused on areas with the highest, temporally and spatially homogeneous use of PPPs.

In Europe, as part of EU regulations, all farmers are required to keep a record of PPP use on each of their fields. These data can be checked during inspection, but are only kept at the farm level and therefore do not provide an overall picture of the PPP use patterns. In Denmark, the farmers’ records are compiled in a national database, with the overall aim of obtaining an accurate picture of PPP use across the country (known as “Pesticide Load”, (Kudsk et al., 2018). According to the Danish Public Administration Act, information from the Danish database is publicly available. It is compulsory for farmers to upload information on crop area and PPP use by crop. Based on the collected information, hot spots of PPP use can be identified where the potential for resistance development can be expected to be high, as illustrated for a group of insecticides, fungicides and herbicides (Figure 6). The detailed information on PPP use available in the database allows for very detailed analyses of PPP use patterns, for instance over time, or by crop and regions (Jørgensen et al., 2019). The information can also be used to find hot spots for potential environmental impact and undesirable PPP effects, such as leaching (Kudsk et al., 2018), or for further investigations on the impact of specific crop rotations on the intensity of PPP use (Jørgensen et al., 2019). To our knowledge, Denmark is so far the only country with such a detailed system for recording the use of PPP.

**Figure 6:**
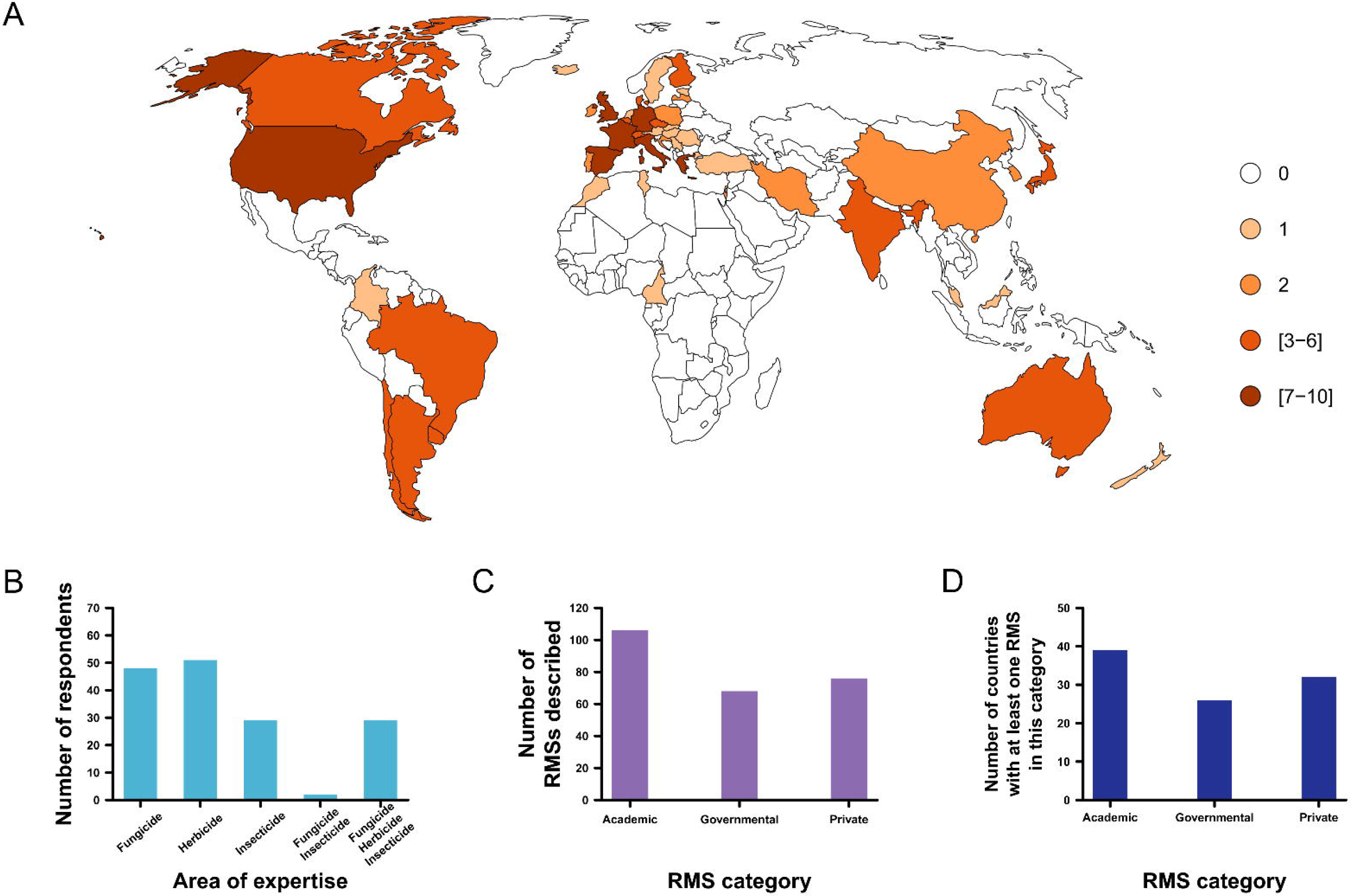
Maps of the intensity of pesticide use (TFI) in Denmark, as an average of four growing seasons (2011-2014). The maps cover three groups of active ingredients: pyrethroids, triazoles, and glyphosate. The maps can provide useful guidance on where the risks for specific resistance problems are higher. Coloring on the maps represents the distribution of calculated values. Blue represents the 8.33% lowest values, and red the 8.33% highest, while the ten color codes in between depict a linear scale with each color code representing 8.33% of the interval between the lowest and highest values.

The Danish Environmental Protection Agency (DEPA) uses data on annual PPP sales and usage. Until now, the data have not been used as background information for resistance monitoring, although this is feasible. Monitoring for PPP resistance in Denmark is mainly organized by academic institutions like Aarhus University, in collaboration with agrochemical companies and advisory services (SEGES), which help to organize the collection of samples across the country. An extensive partnership involving academic, governmental, and private actors addressing fungicide resistance monitoring is led by NorBaRAG, an international Nordic Baltic resistance action group (Kudsk, 2010). This has resulted in several papers providing an overall picture of the resistance situation (Heick et al., 2017).

### Box 3

**Knowing the resistance status as a prerequisite for smart resistance management**

Gathering information on resistance to PPPs requires resources, field-oriented networks of stakeholders, and multi-scale organization within a country. RMS aims can be the detection of emerging resistance and the assessment of the prevalence of resistance in bioagressor populations. This information is crucial for the evaluation of the impact of anti-resistance strategies. In situations where the risk for resistance evolution is high, PPP users are expected to benefit from coordination fostered by sharing RMIPs, but this gain may be at the expense of agrochemical companies that may consequently face lower PPP demand (Lemarié & Marcoul, 2018). This makes information on resistance strategic, and may explain why different RMSs or types of RMSs coexist in the same country.

Resistance is also a hot topic for plant protection, because of growing social pressure towards agriculture that is less dependent on chemical inputs. Consequently, this may also lead to more complex NRMLs including complementary RMSs. In our questionnaire, we identified some initiatives trying to structure resistance monitoring at a national level and centralizing RMIPs (see SI2). As an example, we detail here the case of France, which is best known by the authors. Other countries may of course have equivalent or more comprehensive organizations.

Up to 5 RMSs can be distinguished in France. (1) Monitoring is achieved by agrochemical companies and their contractors (i.e. Private RMSs) to provide data for authorization dossiers, pre- and post-authorization, as required by European and French regulations. This information is used for marketing authorization, issued by the public agency in charge of PPP approvals. (2) Agrochemical companies, but also cooperatives, retailers, and extension services, represented by crop-specialized technical institutes, (i.e. Private RMSs) may also carry out more specific monitoring to accompany PPP development and offer use recommendations throughout the lifespan of the PPP. Such information is used for internal purposes or targeted publication towards stakeholders, and is in part summarized by Resistance Action Committees (RACs) on their websites. (3) Public research institutes also collect and analyze samples, often in collaboration with extension services, for basic research on resistance evolution, management, and mechanisms (i.e. Academic RMSs), which can be used for recommendations on resistance management, together with other stakeholders. (4) The French Ministry of Agriculture funds a national resistance monitoring plan (i.e. a Governmental RMS) aimed at detecting emerging resistances, with situations prioritized by field practitioners and experts. This is legitimated by the Écophyto national regulation that includes the monitoring of non-intended effects of PPP use. Samples are collected via a large range of partners, some being organized in networks. PPP sensitivity assays are achieved by a dedicated public laboratory and by research institutes. Results are used to inform public decision-making. (5) Phytopharmacovigilance (i.e. a Governmental RMS) is organized by the French Ministries of Agriculture, Human Health, and the Environment and request the official declaration of resistance cases by any stakeholders. It relies on the previous RMSs described and encourages all stakeholders to feed back information to identify early signs of resistance. Its data are considered for post-authorization.

Finally, in addition to manufacturer use recommendations, available information from these various RMSs is summarized by representatives of agricultural sectors and general experts in collaborative notes freely available on the internet, in addition to resistance reports (www.r4p-inra.fr). Lists of resistance cases are also regularly updated and reflect agriculture in France today, and its history of PPP use (**Table 1)**. They also orient further research and monitoring on resistance, as well as PPP evaluation.

**Table 1:**
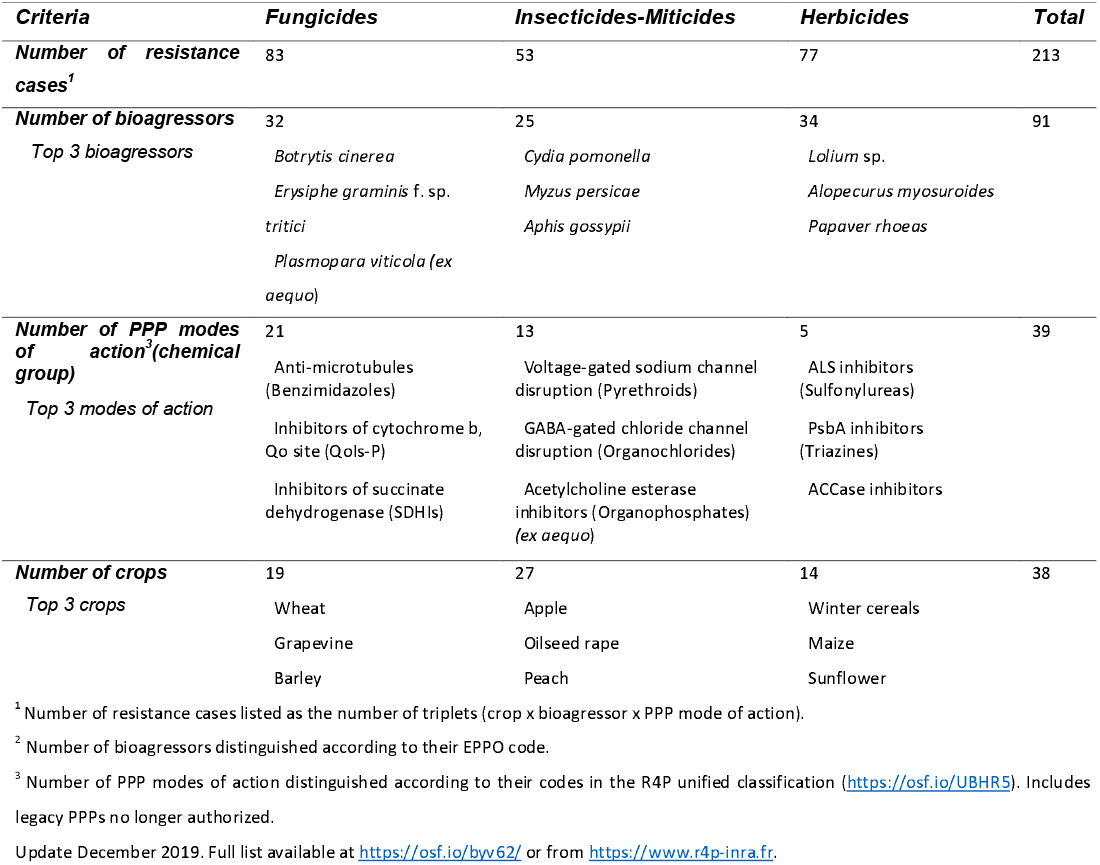
Details on resistance cases detected in France (1978 – 2019).

## FIGURE LEGENDS

**Figure S1**.**1:** Probability of presence of one RMS category in a country depending on the Human Development Index (HDI) of the country. A) Probability of an Academic RMS in the country (non-significant correlation, Z = 0.79, *p* = 0.429), B) Probability of a Governmental RMS in the country (significant correlation, Z = 2.37, *p* = 0.018), C) Probability of a Private RMS in the country (non-significant correlation, Z = 0.32, *p* = 0.746). Logistic regression model prediction (solid line) and 95 % CI (grey band) are presented.

**Figure S1**.**2:** Probability of presence of one type of source for monitoring of PPP use data in a country depending on the Human Development Index (HDI) of the country. A) Probability of PPP use data monitored by PPP sales in the country (significant correlation, Z = 2.34, *p* = 0.019), B) Probability of PPP use data monitored by PPP survey in the country (marginally significant correlation, Z = 1.90, *p* = 0.058). Logistic regression model prediction (solid line) and 95 % CI (grey band) are presented.

## Notes

### Competing Interest Statement

The authors have declared no competing interest.

### Summary of Updates

Results and discussion clarified and less redundant.

