## Supplementary material for "Monitoring systems for resistance to plant protection products across the world: Between redundancy and complementarity": Supporting Information 1-2.pdf

### SI1: The HDI influences NRML diversity and data collection on PPP use

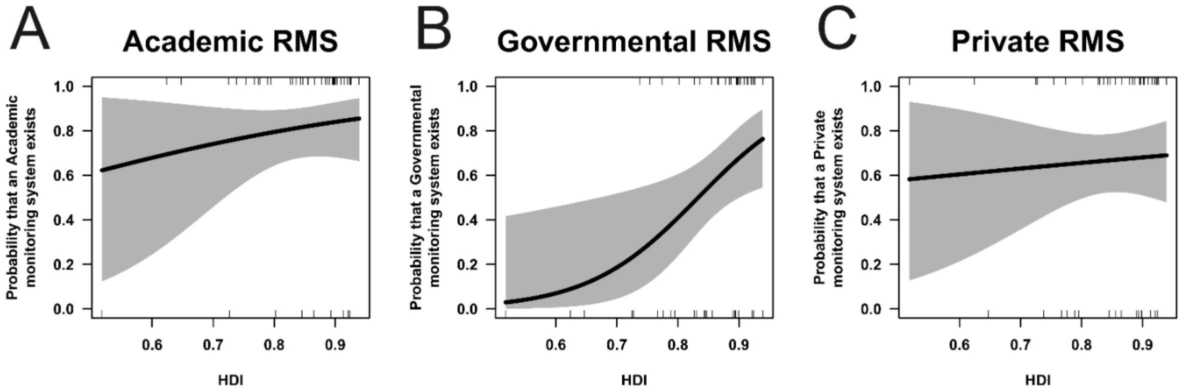


**Figure S1.1:** Probability of presence of one RMS category in a country depending on the Human Development Index (HDI) of the country. A) Probability of an Academic RMS in the country (non-significant correlation, Z = 0.79, *p* = 0.429), B) Probability of a Governmental RMS in the country (significant correlation, Z = 2.37, *p* = 0.018), C) Probability of a Private RMS in the country (non-significant correlation, Z = 0.32, *p* = 0.746). Logistic regression model prediction (solid line) and 95 % CI (grey band) are presented.

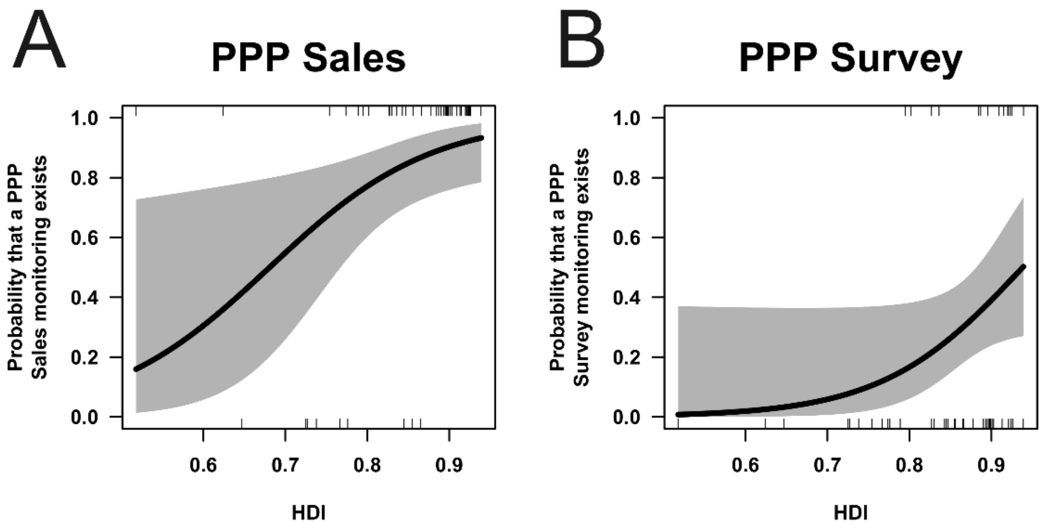


**Figure S1.2:** Probability of presence of one type of source for monitoring of PPP use data in a country depending on the Human Development Index (HDI) of the country. A) Probability of PPP use data monitored by PPP sales in the country (significant correlation, Z = 2.34, $p$ = 0.019), B) Probability of PPP use data monitored by PPP survey in the country (marginally significant correlation, Z = 1.90, $p$ = 0.058). Logistic regression model prediction (solid line) and 95 % CI (grey band) are presented.

**SI2: Internet resources on resistance monitoring groups identified and links used in our study**

- **International Resistance Action Committees**

Fungicide Resistance Action Committee https://www.frac.info/

Herbicide Resistance Action Committee https://www.hracglobal.com/; http://www.weedscience.org; http://wssa.net/wssa/weed/resistance/

Insecticide Resistance Action Committee https://irac-online.org/

Rodenticide Resistance Action Committee https://rrac.info/

- **National resistance monitoring groups mentioning resistance case lists identified in the answers to the questionnaire**

In our questionnaire, we identified some initiatives trying to structure resistance monitoring at a national level and centralizing RMIPs that are worth citing:

- the Research and Reflection Ring on Pesticide Resistance (R4P) Network in France

https://www.r4p-inra.fr

- the Resistance Action Groups (RAG) in the UK

https://ahdb.org.uk/frag

https://ahdb.org.uk/irag

https://ahdb.org.uk/wrag

- the Galanthus resistance database in Greece

http://en.galanthos.gr/

- the Italian Herbicide Resistance Working Group (GIRE)

http://gire.mlib.cnr.it/

- Phytosanitary portal of the Czech Central Institute for Supervising and Testing in Agriculture (UKZUZ)

www.ukzuz.cz/rlportal

- the Japanese Herbicide Resistance Working Group (JHRWG)

http://www.wssj.jp/~hr/index.html

- the New Zealand Committee on Pesticide Resistance (NZCPR)

http://resistance.nzpps.org/

- the Spanish Herbicide Resistance Prevention Comittee (CPRH)

http://semh.net/grupos-de-trabajo/cprh/

- The transnational Nordic Baltic Resistance Action Group (NorBaRAG)

https://projects.au.dk/norbarag/

- **PPP registration authorities across the world cited in Box 1**

US Pesticide Registration Improvement Extension Act PRIA 4: https://www.epa.gov/pesticide-registration

New Zealand: https://www.mpi.govt.nz/processing/agricultural-compounds-and-vet-medicines/acvm-overview/authorisation-of-acvm/

Japan: http://www.acis.famic.go.jp/eng/shinsei/index.htm

FAO Resistance risk assessment guidelines: http://www.fao.org/agriculture/crops/thematic-sitemap/theme/pests/code/list-guide-new/en/

Registration dossiers in Australia: https://apvma.gov.au/node/978

China Regulation on the Administration of Pesticides – Decree 677: https://agrochemical.chemlinked.com/agropedia/overview-chinas-new-pesticide-regulations

https://www.reach24h.com/en/news/industry-news/agrochemical/china-promulgates-new-pesticide-regulation-2.html

- **Legislative texts on PPP regulation**

EU requirement for the recording PPP use at the country level:

Regulation (EC) No 1107/2009 of the European Parliament and of the Council of 21 October 2009 concerning the placing of plant protection products on the market and repealing Council Directives 79/117/EEC and 91/414/EEC article 67 (http://data.europa.eu/eli/reg/2009/1107/oj)

Regulation (EC) No 1185/2009 of the European Parliament and of the Council of 25 November 2009 concerning statistics on pesticides (Text with EEA relevance) (https://eur-lex.europa.eu/eli/reg/2009/1185/oj )

**Directive 2009/128/EC of the European Parliament and of the Council of 21 October 2009 establishing a framework for Community action to achieve the sustainable use of pesticides (Text with EEA relevance) (**http://data.europa.eu/eli/dir/2009/128/oj)

Measures for PPP resistance monitoring in France:

Articles 50 to 54 set out various measures concerning plant protection products, concerning advice, integrated protection, biocontrol, certificates of economy of plant protection products, product traceability, transfer of placing on market authorizations to ANSES (French Agency for Food, Environmental and Occupational Health & Safety) , controls and sanctions on falsification, protection of vulnerable persons, and establishment of a phytopharmacovigilance system.

**Law No. 2014-1170 of 13 October 2014 for the future of agriculture, food and forestry. Title III: Food Policy and Sanitary Performance (**https://www.legifrance.gouv.fr/affichCodeArticle.do?cidTexte=LEGITEXT000006071367&idArticle=LEGIARTI000029581993**)**

- **Database of PPP use in France**

https://ree.developpement-durable.gouv.fr/actualites/article/une-application-de-datavisualisation-des-donnees-d-achats-et-de-ventes-des

**SI3: Online questionnaire used to collect data for this study**
