## Supplementary material for "Monitoring systems for resistance to plant protection products across the world: Between redundancy and complementarity": Supporting Information 1-3.pdf

##### SI1: The HDI influences NRML diversity and data collection on PPP use

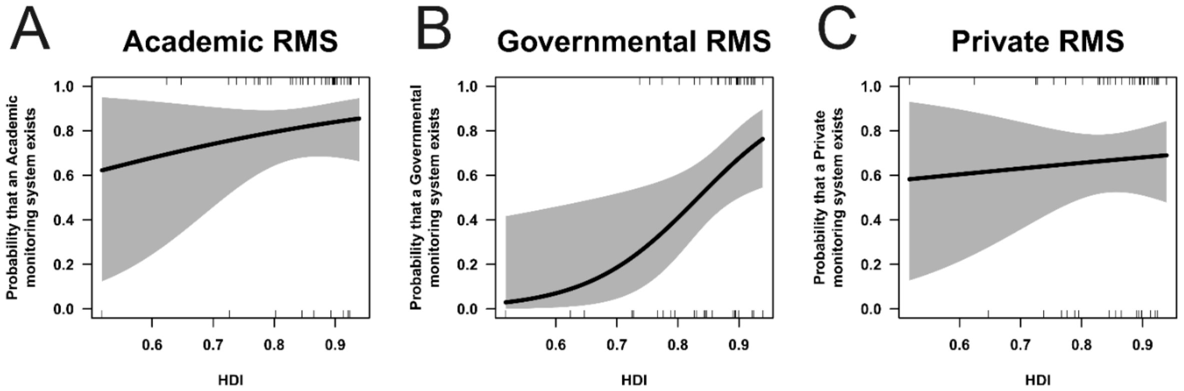

**Figure S1.1:** Probability of presence of one RMS category in a country depending on the Human Development Index (HDI) of the country. A) Probability of an Academic RMS in the country (non-significant correlation, Z = 0.79, $p$ = 0.429), B) Probability of a Governmental RMS in the country (significant correlation, Z = 2.37, $p$ = 0.018), C) Probability of a Private RMS in the country (non-significant correlation, Z = 0.32, $p$ = 0.746). Logistic regression model prediction (solid line) and 95 % CI (grey band) are presented.

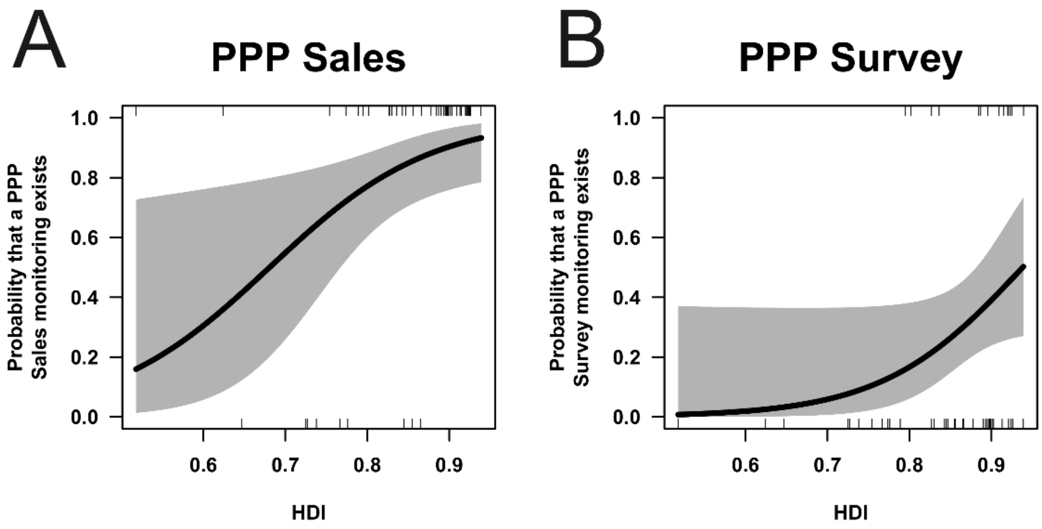

**Figure S1.2:** Probability of presence of one type of source for monitoring of PPP use data in a country depending on the Human Development Index (HDI) of the country. A) Probability of PPP use data monitored by PPP sales in the country (significant correlation, Z = 2.34, $p$ = 0.019), B) Probability of PPP use data monitored by PPP survey in the country (marginally significant correlation, Z = 1.90, $p$ = 0.058). Logistic regression model prediction (solid line) and 95 % CI (grey band) are presented.

**SI2: Internet resources on resistance monitoring groups identified and links used in our study**

- **International Resistance Action Committees**

Fungicide Resistance Action Committee https://www.frac.info/

Herbicide Resistance Action Committee https://www.hracglobal.com/; http://www.weedscience.org; http://wssa.net/wssa/weed/resistance/

Insecticide Resistance Action Committee https://irac-online.org/

Rodenticide Resistance Action Committee https://rrac.info/

- **National resistance monitoring groups mentioning resistance case lists identified in the answers to the questionnaire**

In our questionnaire, we identified some initiatives trying to structure resistance monitoring at a national level and centralizing RMIPs that are worth citing:

- the Research and Reflection Ring on Pesticide Resistance (R4P) Network in France

https://www.r4p-inra.fr

- the Resistance Action Groups (RAG) in the UK

https://ahdb.org.uk/frag

https://ahdb.org.uk/irag

https://ahdb.org.uk/wrag

- the Galanthus resistance database in Greece

http://en.galanthos.gr/

- the Italian Herbicide Resistance Working Group (GIRE)

http://gire.mlib.cnr.it/

- Phytosanitary portal of the Czech Central Institute for Supervising and Testing in Agriculture (UKZUZ)

www.ukzuz.cz/rlportal

- the Japanese Herbicide Resistance Working Group (JHRWG)

http://www.wssj.jp/~hr/index.html

- the New Zealand Committee on Pesticide Resistance (NZCPR)

http://resistance.nzpps.org/

- the Spanish Herbicide Resistance Prevention Comittee (CPRH)

http://semh.net/grupos-de-trabajo/cprh/

- The transnational Nordic Baltic Resistance Action Group (NorBaRAG)

https://projects.au.dk/norbarag/

- **PPP registration authorities across the world cited in Box 1**

US Pesticide Registration Improvement Extension Act PRIA 4: https://www.epa.gov/pesticide-registration

New Zealand: https://www.mpi.govt.nz/processing/agricultural-compounds-and-vet-medicines/acvm-overview/authorisation-of-acvm/

Japan: http://www.acis.famic.go.jp/eng/shinsei/index.htm

FAO Resistance risk assessment guidelines: http://www.fao.org/agriculture/crops/thematic-sitemap/theme/pests/code/list-guide-new/en/

Registration dossiers in Australia: https://apvma.gov.au/node/978

China Regulation on the Administration of Pesticides – Decree 677: https://agrochemical.chemlinked.com/agropedia/overview-chinas-new-pesticide-regulations

https://www.reach24h.com/en/news/industry-news/agrochemical/china-promulgates-new-pesticide-regulation-2.html

- **Legislative texts on PPP regulation**

EU requirement for the recording PPP use at the country level:

Regulation (EC) No 1107/2009 of the European Parliament and of the Council of 21 October 2009 concerning the placing of plant protection products on the market and repealing Council Directives 79/117/EEC and 91/414/EEC article 67 (http://data.europa.eu/eli/reg/2009/1107/oj)

Regulation (EC) No 1185/2009 of the European Parliament and of the Council of 25 November 2009 concerning statistics on pesticides (Text with EEA relevance) (https://eur-lex.europa.eu/eli/reg/2009/1185/oj )

**Directive 2009/128/EC of the European Parliament and of the Council of 21 October 2009 establishing a framework for Community action to achieve the sustainable use of pesticides (Text with EEA relevance) (**http://data.europa.eu/eli/dir/2009/128/oj)

Measures for PPP resistance monitoring in France:

Articles 50 to 54 set out various measures concerning plant protection products, concerning advice, integrated protection, biocontrol, certificates of economy of plant protection products, product traceability, transfer of placing on market authorizations to ANSES (French Agency for Food, Environmental and Occupational Health & Safety) , controls and sanctions on falsification, protection of vulnerable persons, and establishment of a phytopharmacovigilance system.

**Law No. 2014-1170 of 13 October 2014 for the future of agriculture, food and forestry. Title III: Food Policy and Sanitary Performance (**https://www.legifrance.gouv.fr/affichCodeArticle.do?cidTexte=LEGITEXT000006071367&idArticle=LEGIARTI000029581993**)**

- **Database of PPP use in France**

https://ree.developpement-durable.gouv.fr/actualites/article/une-application-de-datavisualisation-des-donnees-d-achats-et-de-ventes-des

**SI3: Online questionnaire used to collect data for this study**

### Plant protection product resistances monitoring systems across the world

#### Introduction 1/2

The aim of this survey is to provide an overview on:

- how resistances to plant protection products are monitored in as many countries as possible worldwide,
- what type of data are available in each country surveyed.

The first part of the survey will give you the opportunity to describe how resistance to plant protection products is monitored in your country.

The second part of the survey is about the available data regarding pesticide use in your country. Indeed, knowing the use of plant protection products can improve the efficacy of  plant protection product resistance monitoring.

The results and conclusion drawn from this survey will be part of a review paper on pesticide resistance monitoring. The published article (pdf) will be send to all the survey respondents in order to thank them.

### Plant protection product resistances monitoring systems across the world

#### Introduction 2/2

Thank you in advance for taking some time (~30 min) to complete this survey. You can refer to the pdf version of the survey to identify the documents needed to answers the questions. **Please complete the survey in one session as the data will be lost if you leave the survey before submitting your answers.**

You can contact us for any questions regarding this project or if you have problems completing the survey:.

In all the survey, **PPP** means Plant Protection Product(s).

You may answer on what you know or/and what you do. Please use the answer **"do not know"** if you do not know the answers.

Thank you for your participation!

Previous Next

### General information

**Thank you to indicate your contact details :**

| |
|---|
| Name |
| Country |
| Organisation |
| Function |
| e-mail |

**Are you expert on a specific group of PPP? Which one?**
*In all the survey,* ***PPP*** *means Plant Protection Product(s).*

- ☐ insecticide
- ☐ herbicide
- ☐ fungicide
- ☐ rodenticide
- ☐ generalist

Previous Next

### Monitorings of resistance to PPP 1/2

#### How resistances to PPP are monitored in your country?

There are different kind of organisations:

• official and nationwide monitorings, usually supported by the public administration, many stakeholders can participate. It can include post-registration monitorings.

• private monitorings, conducted either by companies, distributors or technical institutes independantly from the official and nationwide monitoring. It can include private post-registration monitorings.

• academic research projects on resistance to PPP, independant from the to other kind of monitorings.

In one country, different monitorings can be conducted.

**Please indicate which kind of monitorings are conducted in your country. Then some questions will give your the opportunity to give details on each of monitoring who selected.**

**Are there a national/federal and official monitorings of resistances to PPP in your country?**

○ yes ○ no ○ do not know

**Are there private monitorings of resistances to PPP in your country?**

○ yes ○ no ○ do not know

**Are there some academic research projects on resistances to PPP in your country?**

○ yes ○ no ○ do not know

*These projects must not be a part of the official and nationwide monitoring or of a private monitoring.*

Previous Next

### Objectives and scope of the nationwide and official monitoring

Previously, you answered that there are national/federal and official monitorings of resistances to PPP in your country. The following questions focus on this(these) national/federal and official monitoring(s) of resistances to PPP.

**What are the objectives of this monitoring?**

- ○ emergence
- ○ frequency of existing resistances
- ○ both
- ○ do not know

**How many resistance themes are studied in your country (theme = crop * active substance * pest)?**

**Details on the resistance themes**

**Who participate to this choice?**

- ☐ Public administration
- ☐ Academics
- ☐ Companies
- ☐ do not know
- ☐ other

Specify if you answered "other"

**Do the monitoring results have an impact on the registration procedure of PPP in your country?**

- ○ yes
- ○ no
- ○ do not know

Previous | Next

### Institutional organisation of the nationwide and official monitoring

*A monitoring can be organised at different scales. It depends usually on the administrative organisation of your country (federal state or not).*

*In the following questions, we distinguish two scales :*

- *national, ie. centralisation at the national level or the highest administrative level you are able to describe*
- *regional, ie. a lower administrative level than the national one : state in a federal state, region, ...*

*Please feel free to answer on what you know.*

**Please give the scales you will describe :**

| |
|---|
| national (the highest administrative level you are able to describe) |
| regional (a lower administrative level than the national one) |

Previous Next

#### Who manages the nationwide and official monitoring of resistances at the national scale?

**Leader(s).** *Please give name, organisation, e-mail.*

**Participants**. *Give an example of the stakeholders involved.*

**Who funds the monitoring of resistances at the national scale?**

- ☐ Public administration
- ☐ Academics
- ☐ Companies
- ☐ do not know
- ☐ other

Specify if you answered "other"

Previous Next

#### Who manages the nationwide and official monitoring of resistances at the regional scale?

**Leader(s) (if known).** *Please give name, organisation, e-mail.*

**Participants.** *Give an example of the stakeholders involved.*

**Are all the regions of your country involved similarly in the surveillance of resistances?**

- yes
- do not know
- no

details

**Who collects the samples in the field?**

- officials
- advisers
- local or regional agricultural administration
- distributors
- academics
- companies
- do not know
- other

Specify if you answered "other"

**How many samples are yearly collected (on average for each resistance themes - crop * active substance * pest)?** *A sample corresponds to a field / a location.*

Previous Next

### Cooperation with other countries

**Does your country cooperate with others to monitor resistances?**

○ yes ○ no ○ do not know

**Please give your partner countries:**

**Please give an e-mail address of your partner(s) abroad:**

Previous Next

### Laboratory

**What is the status of the laboratories involved in the nationwide and official monitoring?**

- ☐ public laboratory
- ☐ academic laboratory
- ☐ company laboratory
- ☐ service laboratory (other than company)
- ☐ do not know
- ☐ other

Specify if you answered "other" ____________________

**Which analysis techniques are used?**

- ☐ biotest
- ☐ biomolecular
- ☐ biochemical
- ☐ do not know

Previous | Next

### Monitoring tools and procedure

**Is there an official (formalised) sampling protocol?**

- ( ) yes
- ( ) no
- ( ) do not know

**How are sampling locations chosen?**

- [ ] randomly
- [ ] based on selection pressure (ie PPP use)
- [ ] loss of efficacy of the treatment
- [ ] do not know
- [ ] other

details:

**What kind of surveillance is made?**

- ( ) passive (spontaneous report of resistance suspicion from the field)
- ( ) active (scheduled sampling campaign)
- ( ) both
- ( ) do not know

Previous Next

### Data management for the nationwide and official monitoring

**Which data are collected on the field?**

- ☐ crop
- ☐ GPS coordinates
- ☐ plant protection strategy used
- ☐ disease intensity
- ☐ do not know
- ☐ other

if 'other' precise:

**How are they collected?**

- ☐ paper form
- ☐ electronic form
- ☐ do not know

**Is there a national database?**

- ○ yes
- ○ no
- ○ do not know

**Who analyzes and interprets the data?**

- ☐ officials
- ☐ Academics
- ☐ Companies
- ☐ Advisers
- ☐ Distributors
- ☐ do not know
- ☐ other

if 'other', precise:

Previous | Next

### Communication

**Are the results of the monitoring published regularly?** *Several choices authorized, eg in case the frequency of publication of the different theme varies.*

- ☐ during the cropping season
- ☐ yearly
- ☐ less frequently
- ☐ not published (internal use only)
- ☐ do not know
- ☐ other

details:

**How are they diffused?**

- ☐ annual review meeting
- ☐ scientific publications
- ☐ technical publications
- ☐ news letters
- ☐ EPPO
- ☐ dedicated website
- ☐ do not know
- ☐ other

Specify if you answered "other"

**if website, please give the address**

**In which language?**

**Who are targeted by the publications?**

- ☐ farmers
- ☐ scientists
- ☐ regulatory authorities
- ☐ do not know
- ☐ other

details

Previous Next

### Objectives and scope of private monitorings

Previously, you answered that there are private monitorings of resistances to PPP in your country. The following questions focus on this(these) private monitoring(s) of resistances to PPP.

**What are the objectives of the private monitorings?**

- ◯ emergence
- ◯ frequency of existing resistances
- ◯ both
- ◯ do not know

**How many resistance themes are studied in your country (theme = crop * active substance * pest)?**

**Details on the resistance themes**

**Who participate to this choice?**

- ☐ Public administration
- ☐ Academics
- ☐ Companies
- ☐ do not know
- ☐ other

Specify if you answered "other"

**Do the monitoring results have an impact on the registration procedure of PPP in your country?**

- ◯ yes
- ◯ no
- ◯ do not know

Previous | Next

### Institutional organisation of the private monitorings

*A monitoring can be organised at different scales. It depends usually on the administrative organisation of your country (federal state or not).*

*In the following questions, we distinguish two scales :*

- *national, ie. centralisation at the national level or the highest administrative level you are able to describe*
- *regional, ie. a lower administrative level than the national one : state in a federal state, region, ...*

*Please feel free to answer on what you know.*

**Please give the scales you will describe :**

| |
|---|
| national (the highest administrative level you are able to describe) |
| regional (a lower administrative level than the national one) |

**Are these private monitorings :**

- ☐ international
- ☐ national
- ☐ regional
- ☐ do not know

Previous Next

#### Who manages the private monitorings of resistances at the <u>national</u> scale?

**Leader(s).** *Please give name, organisation, e-mail.*

**Participants.** *Give an example of the stakeholders involved.*

**How are these private monitorings funded? Who funds these monitorings?**

Previous Next

#### Who manages the private monitoring of resistances at the regional scale?

**Leader(s) (if known).** *Please give name, organisation, e-mail.*

**Participants.** *Give an example of the stakeholders involved.*

**Are the private monitorings done similarly in all the regions of your country?**

yes    do not know    no

details

**How many samples are yearly collected (on average for each resistance themes - crop * active substance * pest)?** *A sample corresponds to a field / a location.*

Previous    Next

### Laboratory for private monitorings

**What is the status of the laboratories involved in the private monitorings?**

- ☐ public laboratory
- ☐ academic laboratory
- ☐ company laboratory
- ☐ service laboratory (other than company)
- ☐ do not know
- ☐ other

Specify if you answered "other" ____________________

**Which analysis techniques are used?**

- ☐ biotest
- ☐ biomolecular
- ☐ biochemical
- ☐ do not know

Previous | Next

### Monitoring tools and procedure for private monitorings

**Is there an official (formalised) sampling protocol?**

- yes
- no
- do not know

**How are sampling locations chosen?**

- randomly
- based on selection pressure (ie PPP use)
- loss of efficacy of the treatment
- do not know
- other

details:

**What kind of surveillance is made?**

- passive (spontaneous report of resistance suspicion from the field)
- active (scheduled sampling campaign)
- both
- do not know

### Data management for private monitorings

**Which data are collected on the field?**

- [ ] crop
- [ ] GPS coordinates
- [ ] plant protection strategy used
- [ ] disease intensity
- [ ] do not know
- [ ] other

if 'other' precise:

**How are they collected?**

- [ ] paper form
- [ ] electronic form
- [ ] do not know

**Who analyzes and interprets the data?**

- [ ] Officials
- [ ] Academics
- [ ] Companies
- [ ] Advisers
- [ ] Distributors
- [ ] do not know
- [ ] other

if 'other', precise:

**Are these data effectively open to other institutions for analysis?**

- ( ) yes
- ( ) no
- ( ) do not know

### Communication in private monitorings

**Are the results of the monitoring published regularly?** *Several choices authorized, eg in case the frequency of publication of the different theme varies.*

- [ ] during the cropping season
- [ ] yearly
- [ ] less frequently
- [ ] not published (internal use only)
- [ ] do not know
- [ ] other

details:

**How are they diffused?**

- [ ] annual review meeting
- [ ] scientific publications
- [ ] technical publications
- [ ] news letters
- [ ] EPPO
- [ ] dedicated website
- [ ] do not know
- [ ] other

Specify if you answered "other"

**if website, please give the address**

**In which language?**

**Who are targeted by the publications?**

- [ ] farmers
- [ ] scientists
- [ ] regulatory authorities
- [ ] do not know
- [ ] other

details:

Previous Next

### Objectives and scope of monitorings (academic research projects)

Previously, you answered that there are academic research projects on resistances to PPP in your country. The following questions focus on this(these) academic research projects.

**What are the objectives of the monitorings done in research projects?**

- ○ emergence
- ○ frequency of existing resistances
- ○ both
- ○ do not know

**How many resistance themes are studied in your country (theme = crop * active substance * pest) in research projects?**

**Details on the resistance themes**

**Who participate to this choice?**

- ☐ Public administration
- ☐ Academics
- ☐ Companies
- ☐ do not know
- ☐ other

Specify if you answered "other"

**Do the monitoring results have an impact on the registration procedure of PPP in your country?**

- ○ yes
- ○ no
- ○ do not know

### Institutional organisation of academic research projects

**Leader(s).** *Please give name, organisation, e-mail.*

**Participants.** *Give an example of the stakeholders involved.*

**How are these research projects funded? Who funds these projects?**

**What is the usual geographical scale of the research projects on resistance to PPP?**

☐ nationwide ☐ do not know ☐ regional

details

**How many samples are yearly collected (on average for each resistance themes - crop * active substance * pest)?** *A sample corresponds to a field / a location.*

Previous Next

### Laboratory

**What is the status of the laboratories involved in the academic research projects?**

- ☐ public laboratory
- ☐ academic laboratory
- ☐ company laboratory
- ☐ service laboratory (other than company)
- ☐ do not know
- ☐ other

Specify if you answered "other"

**Which analysis techniques are used?**

- ☐ biotest
- ☐ biomolecular
- ☐ biochemical
- ☐ do not know

### Monitoring tools and procedure

**Is there an official (formalised) sampling protocol?**

- ○ yes
- ○ no
- ○ do not know

**How are sampling locations chosen?**

- ☐ randomly
- ☐ based on selection pressure (ie PPP use)
- ☐ loss of efficacy of the treatment
- ☐ do not know
- ☐ other

details:

**What kind of surveillance is made?**

- ○ passive (spontaneous report of resistance suspicion from the field)
- ○ active (scheduled sampling campaign)
- ○ both
- ○ do not know

### Data management for academic research projects

**Which data are collected on the field?**

- ☐ crop
- ☐ GPS coordinates
- ☐ plant protection strategy used
- ☐ disease intensity
- ☐ do not know
- ☐ other

if 'other' precise:

**How are they collected?**

- ☐ paper form
- ☐ electronic form
- ☐ do not know

**Is there a national database?**

- ○ yes
- ○ no
- ○ do not know

**Who analyzes and interprets the data?**

- ☐ Officials
- ☐ Academics
- ☐ Companies
- ☐ Advisers
- ☐ Distributors
- ☐ do not know
- ☐ other

if 'other', precise:

### Communication in academic research projects

**Are the results of the monitoring published regularly?** *Several choices authorized, eg in case the frequency of publication of the different theme varies.*

- ☐ during the cropping season
- ☐ yearly
- ☐ less frequently
- ☐ not published (internal use only)
- ☐ do not know
- ☐ other

details:

**How are they diffused?**

- ☐ annual review meeting
- ☐ scientific publications
- ☐ technical publications
- ☐ news letters
- ☐ EPPO
- ☐ dedicated website
- ☐ do not know
- ☐ other

Specify if you answered "other"

**if website, please give the address**

**In which language?**

**Who are targeted by the publications?**

- ☐ farmers
- ☐ scientists
- ☐ regulatory authorities
- ☐ do not know
- ☐ other

details:

Previous | Next

### Monitorings of resistance to PPP 2/2

**Do you know other monitorings of resistance to PPP in your country you did not mention in the previous questions?**

- no
- yes

details:

**If there are different monitorings in your country (official, private, research projects), are all the results compiled?**

- yes
- no
- do not know

**Are there published lists of resistance cases updated regularly in your country?**

- yes
- no
- do not know

**Comments on this part on the different monitorings in your country :**

Previous

Next

### Consequences on the PPP registration and regulation

**Do public authorities impose changes or recommendations on PPP use when new resistances arise?**

- ○ yes
- ○ no
- ○ do not know

**What kind of changes or recommendations?**

- ☐ active substance substitution
- ☐ active substance limitation (lower number of applications per year)
- ☐ PPP removal in specific areas
- ☐ used of the mode of action in sequence with others
- ☐ do not know
- ☐ other

if 'other', precise :

**At which scale?**

- ☐ national
- ☐ regional
- ☐ landscape
- ☐ field
- ☐ do not know
- ☐ other

if 'other', precise :

#### Literature

**If documents about PPP resistance monitoring in your country are available, we would be glad to have access to them.**

**Please fill free to upload them using the following link.**

Ajouter un document

**Or provide the URL to have access to them**

Previous Next

### Knowledge of the PPP use throughout the country

**Are data on PPP use collected in your country?**

- [ ] through sales figures
- [ ] no
- [ ] do not know
- [ ] other

other

Previous Next

### Data collection through PPP sales figures

**Since when are they collected?**

*Please give a year ####*

**Are they collected yearly?**

- yes
- no
- do not know

**How are they collected?**

- surveys
- automatically by the distributor
- do not know
- other

details:

**What is the smallest geographical scale of these data?**

- country
- region
- sale location
- postal code of the user
- parish
- field
- do not know
- other

if 'other', please precise:

**Are these data available and used in some way to monitor the resistance to PPP?**

- no
- do not know
- yes

if 'yes', precise:

Previous | Next

### Other method of data collection on PPP use 1/2

**Are these data on PPP use (<u>other than PPP sales figures</u>) about :**

- ☐ mode of action
- ☐ substance
- ☐ commercial product
- ☐ do not know

**How is measured the PPP quantity?**

- ☐ treatment frequency index
- ☐ tons
- ☐ do not know
- ☐ other

if 'other', please precise:

**Since when are they collected?**

*Please give a year ####*

**Are they collected yearly?**

- ○ yes
- ○ no
- ○ do not know

**How are they collected?**

- ☐ surveys
- ☐ automatically by the distributor
- ☐ do not know
- ☐ other

details:

Previous    Next

#### Other method of data collection on PPP use 2/2

**What is the smallest geographical scale of these data?**

- ○ country
- ○ region
- ○ sale location
- ○ postal code of the user
- ○ parish
- ○ field
- ○ do not know
- ○ other

if 'other', please precise:

**Are these data available and used in some way to monitor the resistance to PPP?**

- ○ no
- ○ do not know
- ○ yes

if 'yes', precise:

### General comments

**General comments on PPP resistances monitoring in your country**

**If you know people in your country or other countries who could answer this survey and have a different expertise than you (on insecticide, herbicide, fungicide), please enter their e-mail address here.**

Previous Next

### Thank you

The results and conclusion drawn from this survey will be part of a review paper on pesticide resistance monitoring. The published article (pdf) will be send to all the survey respondents, in order to thank them.

We plan to attach a list providing the names of the respondents to the article.

**Do you prefer to remain anonymous ?**

○ yes ○ no

Thank you very much for your time and cooperation.

We are available for any questions or discussion regarding this survey.

**Contacts:**

Previous Save
