## Supplementary figures and images for "Monitoring systems for resistance to plant protection products across the world: Between redundancy and complementarity"

### FigureS1.1.png

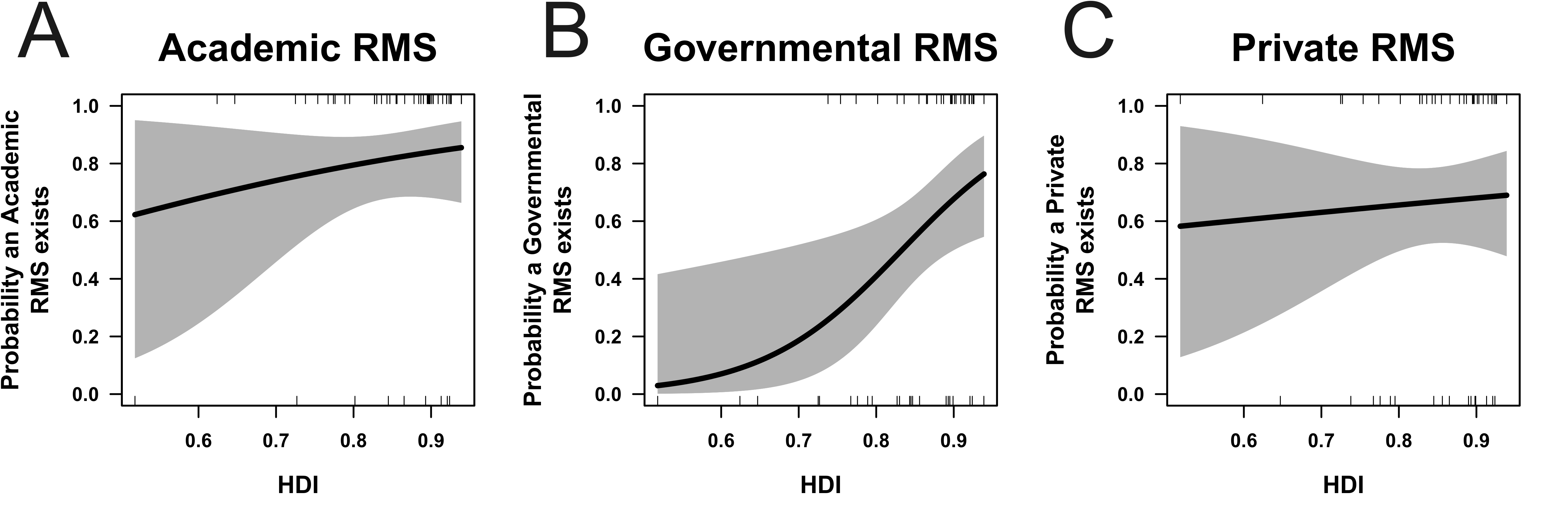

### FigureS1.2.png

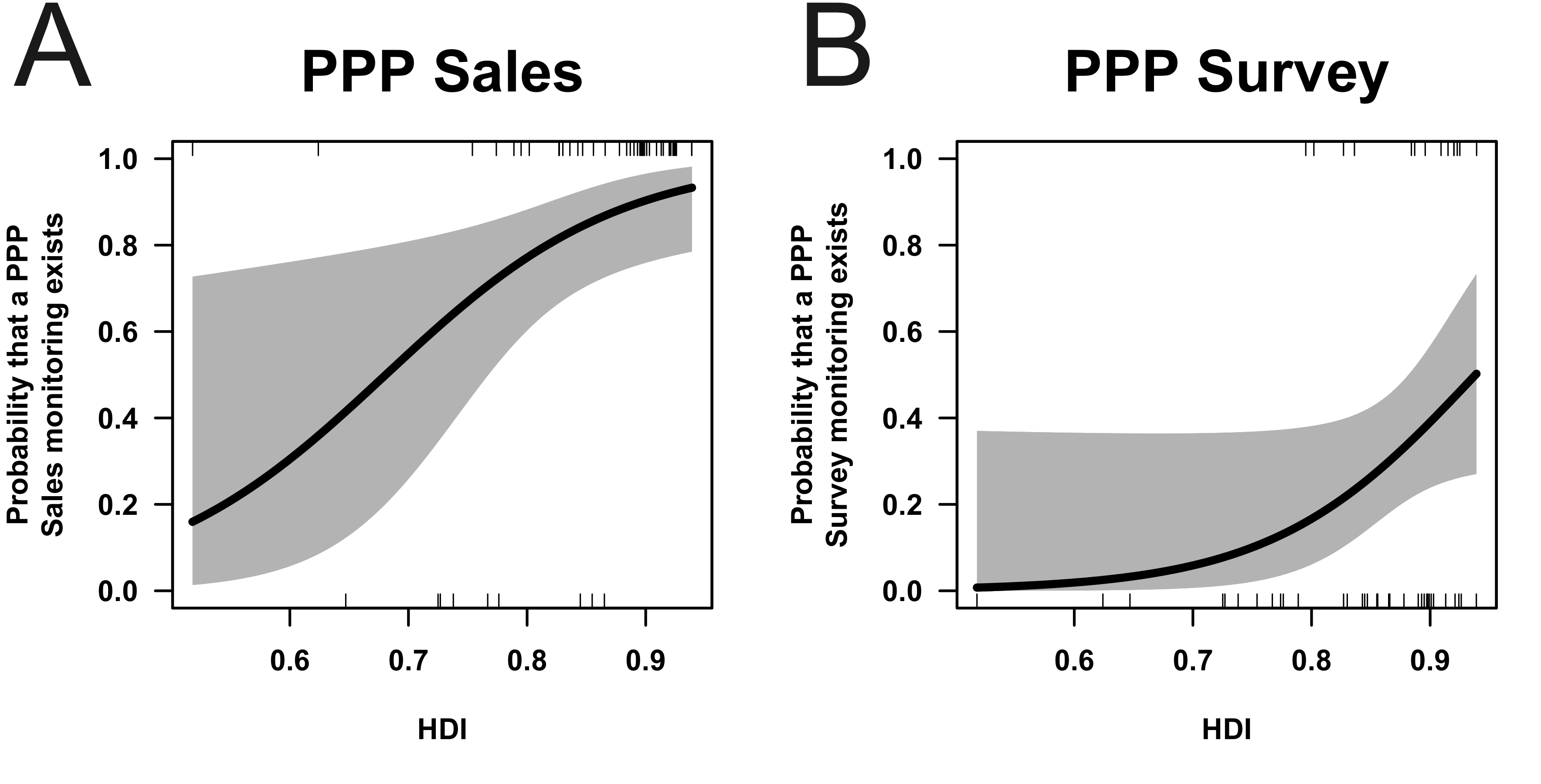
